# Octopamine signals coordinate the spatial pattern of presynaptic machineries within individual *Drosophila* mushroom body neurons

**DOI:** 10.1101/2024.12.02.626512

**Authors:** Hongyang Wu, Sayaka Eno, Kyoko Jinnai, Yoh Maekawa, Kokoro Saito, Ayako Abe, Darren W Williams, Nobuhiro Yamagata, Shu Kondo, Hiromu Tanimoto

**Author notes:** Correspondence (H.T.). These authors contributed equally (Hongyang Wu, Sayaka Eno).

## Abstract

Neurons need to adjust synaptic output according to the targets. However, the target-specific synaptic structures within individual neurons in the central nervous system remains unresolved. Applying the CRISPR/Cas9-mediated split-GFP tagging, we visualized the endogenous active zone scaffold protein, Bruchpilot (Brp), in specific cells. This technology enabled the spatial characterization of presynaptic machineries only within the Kenyon cells (KCs) of the *Drosophila* mushroom bodies. We found the patterned accumulation of Brp among the compartments of axon terminals, where a KC synapses onto different postsynaptic neurons. Mechanistically, the localized octopaminergic modulation along γ KC terminals regulate this compartmental Brp heterogeneity via Octβ2R and cAMP signaling. We further found that acute food deprivation reorganizes this spatial pattern in an octopamine-dependent manner. Such coordinated regulation of local synaptic machineries thus explains how the mushroom bodies integrate changing physiological states. This subcellular information processing represents an elegant solution to expand computational capacity of the circuit.

## INTRODUCTION

The molecular composition of chemical synapses determines the function of neurons and therefore exhibits a remarkable diversity among cell types ^1,2^. As demonstrated by studies on the motor neuron, presynaptic molecular assemblies can be highly heterogeneous even within a single cell ^3,4^. Critically, the difference of presynaptic structures may result in varying synaptic functions at the single active zone (AZ) level, such as release probability ^5–7^. Given that a single neuron in the central nervous system (CNS) typically synapses onto multiple target cells, the spatial adaptation of output machineries could differentiate the activities of post-synaptic cells. While such intracellular synaptic heterogeneity influences complex computation of the circuit in the CNS, the target-specific AZ regulation is scarcely studied.

To this end, neurons projecting to the *Drosophila* mushroom bodies (MBs) can serve as an excellent model system. Kenyon cells (KCs), the major MB intrinsic neurons, synapse onto five sets of post-synaptic partners in spatially segregated compartments ^8,9^. Postsynaptic MB output neurons (MBONs) have distinct and compartmentalized dendrites within the MB lobes ^9^, and their activities collectively determine the MB output ^10–14^. The compartmental distinctions in presynaptic assemblies within individual KCs are likely to be critical in controlling MB-guided behaviors. However, it is challenging to characterize AZs only in KCs without 3D electron microscopy, because the synapse density in MB lobes is among the highest in the fly brain ^15,16^.

We here aim to characterize the compartmental distinctions of presynaptic structures of KCs by visualizing endogenous Brp in a cell-type specific manner. *Drosophila* ELKS/CAST/ERC family member Brp is a major scaffold protein forming the T-shape electron-dense projections decorating AZs ^17–20^. Brp clusters determine molecular assemblies at AZs such as the accumulation of calcium channels and synaptic vesicles ^17,21^. Therefore, Brp structures serve as a proxy for estimating the synapse function. To label endogenous pre-synaptic proteins in designated cell types ^22,23^, we took advantage of the CRISPR/Cas9-mediated split-GFP tagging strategy to target Brp in the dense network of MBs ^24,25^.

Profiling the character of individual Brp clusters only within KCs revealed the compartmental Brp accumulation pattern along the MB lobes. We further report the state-dependent remodeling of this intracellularly organized AZ structure and the regulatory mechanisms.

## RESULTS

### Cell-type specific visualization of endogenous active zone scaffold proteins Brp

Chemical tagging using Brp::SNAP ^26^ suggests endogenous Brp localizes heterogeneously within the MB (Fig. S1), which is constituted by pre-synapses of various cell types (Fig. S2). To visualize endogenous Brp only in designated cell types, we inserted the GFP_11_ fragment (the eleventh β-strand of the super-folder GFP) just prior to the stop codon of *brp* using CRISPR/Cas9. The self-assembly and reconstitution of GFP is induced by expressing the GFP_1-10_ fragments using the GAL4/UAS system (Fig. 1A).

**Figure 1.**
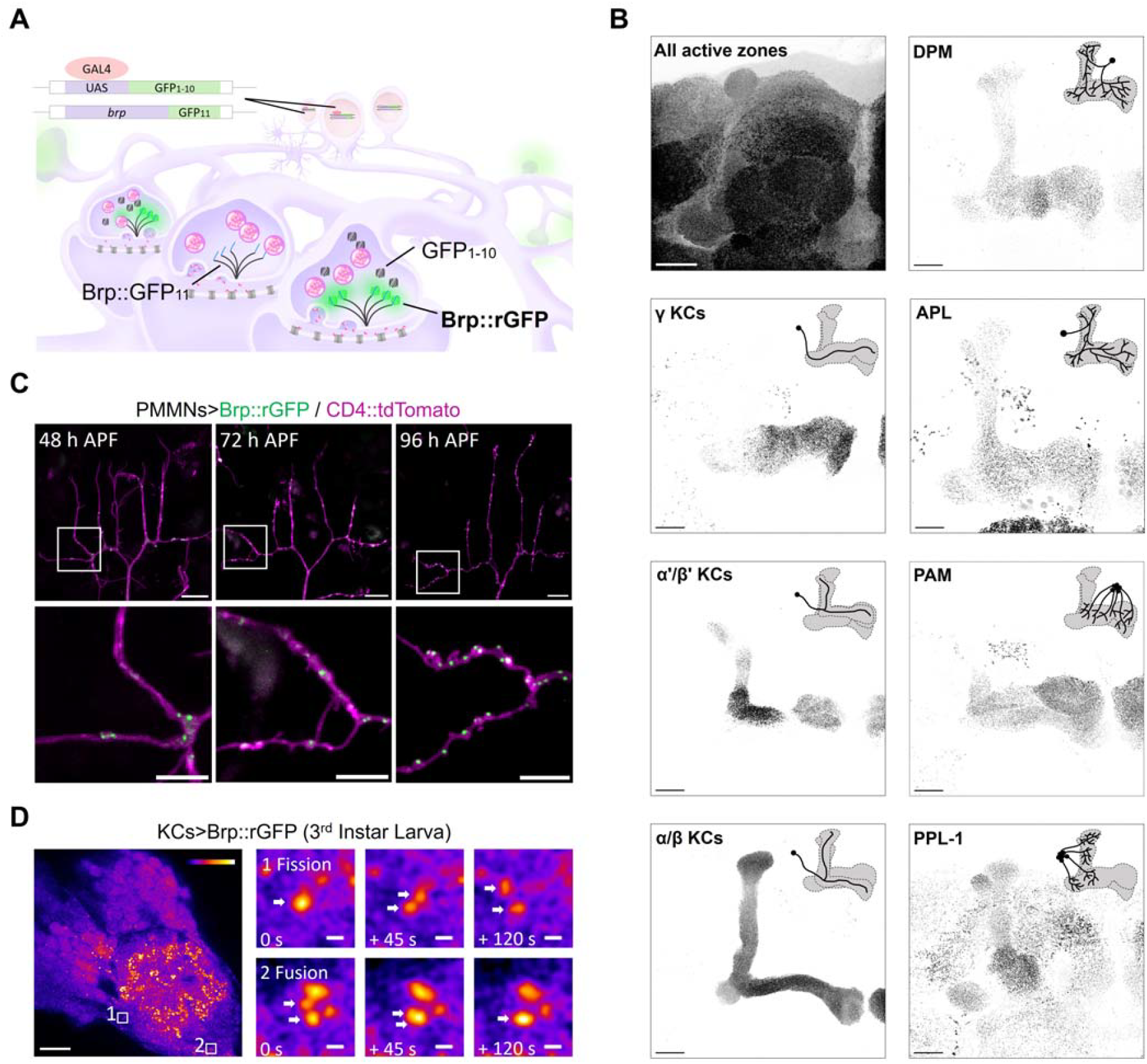
Cell-type specific dissection of the endogenous active zone scaffold. (**A**) Schematic of split-GFP tagging system. GFP_11_ was inserted prior to the stop codon of *brp*, while GFP_1-10_ was expressed in specific cell types using the GAL4-UAS system. GFP_11_ and GFP_1-10_ reconstitute to emit fluorescence. (**B**) Visualization of Brp::rGFP in a cell-type specific manner. Immunostaining using antibody nc82 which labels all Brp in the brain, as a comparison to Brp::rGFP. Diagrams in each panel show the innervation pattern of neurons in the MB. GAL4 lines used: γ KCs (MB009B), α/β KCs (MB008B), α’/β’ KCs (MB370B), PPL-1 (TH-GAL4), PAM (R58E02), DPM (VT64246), APL (GH146). Scale bar, 20 μm. (**C**) Long-term live-imaging of Brp::rGFP (green) in a growing motor neuron (magenta, CD4::tdTomato) from 48 to 96 hours APF. OK371-GAL4 was used to express GFP_1-10_ and CD4::tdTomato. White boxes indicate zoomed-in areas shown in the lower panels. Scale bars: upper panels, 10 μm; lower panels, 5 μm. (**D**) Ex vivo live-imaging of Brp::rGFP in 3^rd^ instar larval KCs. GFP_1-10_ was expressed using R13F02-GAL4. The left panel shows the MB calyx region. White boxes indicate zoomed-in areas shown on the right panels. Box 1 and 2 demonstrate Brp::rGFP fission and fusion event. Whites arrows indicate Brp::rGFP clusters undergoing fission or fusion. Scale bars: left panel, 10 μm; right panels, 200 nm.

We validated this system with GAL4 driver lines that label 7 different MB-projecting neuronal populations: 3 KC subtypes, 2 dopamine neuron clusters (PPL1 and PAM) and 2 single interneurons (DPM and APL neurons) ^9,27^. Confocal microscopy detected reconstituted GFP (rGFP) signals only at restricted areas, in contrast to the dense signals in Brp::SNAP or Anti-Brp immunostaining (Fig. 1B, Fig. S1). The Brp::rGFP signal appeared as individual puncta and matched the areas of axon terminals of the cell types (Fig. 1B). Especially, Brp can be labeled at single-cell resolution for the APL and DPM neuron ^8,28,29^. Strikingly, the Brp::rGFP signal intensity appeared to be heterogenous along the axon terminals of KCs (Fig. 1B). Since individual KCs arborize onto the entire lobe structure, these results suggest distinct accumulations of Brp intracellularly.

To test whether Brp::rGFP labels dynamic synaptic structures, we benchmarked its performance with different live-imaging contexts. The axon arborizations of pleural muscle motor neurons (PM-Mns) can be imaged on the abdominal body wall during pupal development ^30^. Live-imaging Brp::rGFP in the PM-Mn revealed the formation of individual clusters at 48 hours after puparium formation and the increasing density in the same branch following two days (Fig. 1C). We further visualized Brp::rGFP dynamics in the ex-vivo preparation of the larval brain, and found multiple fusion and fission events of Brp clusters in the MB calyx (Fig. 1D). These results are consistent with the AZ assembly through liquid-liquid phase separation ^31^ and validate the method to visualize the dynamics of endogenous Brp structures.

### Intracellular active zone heterogeneity among MB compartments

Brp::rGFP accumulation was heterogeneous among KC compartments, especially in γ KCs (Fig. 1B, 2A, Fig. S3). To characterize this presynaptic Brp heterogeneity, an image-processing pipeline was developed to better quantify the signal intensity of individual Brp::rGFP puncta. Optimizing the parameter combination of 3D image deconvolution ^32,33^ and 3D spot segmentation enabled us to resolve Brp::rGFP clusters that represent single AZs using conventional confocal microscopy ^29^. By sampling a volume of each compartment, we quantified the median of integrated Brp::rGFP intensity of clusters at the AZs. Therefore, the quantification is independent of the AZ density or the sample volume. The analysis revealed a striking compartmental heterogeneity of AZs (Fig. 2B). The median Brp::rGFP intensity of clusters in the γ5 compartment was nearly three times as intense as that in γ1. These results suggest that individual KCs tune their presynaptic structures according to compartments (Fig. 2C).

**Figure 2.**
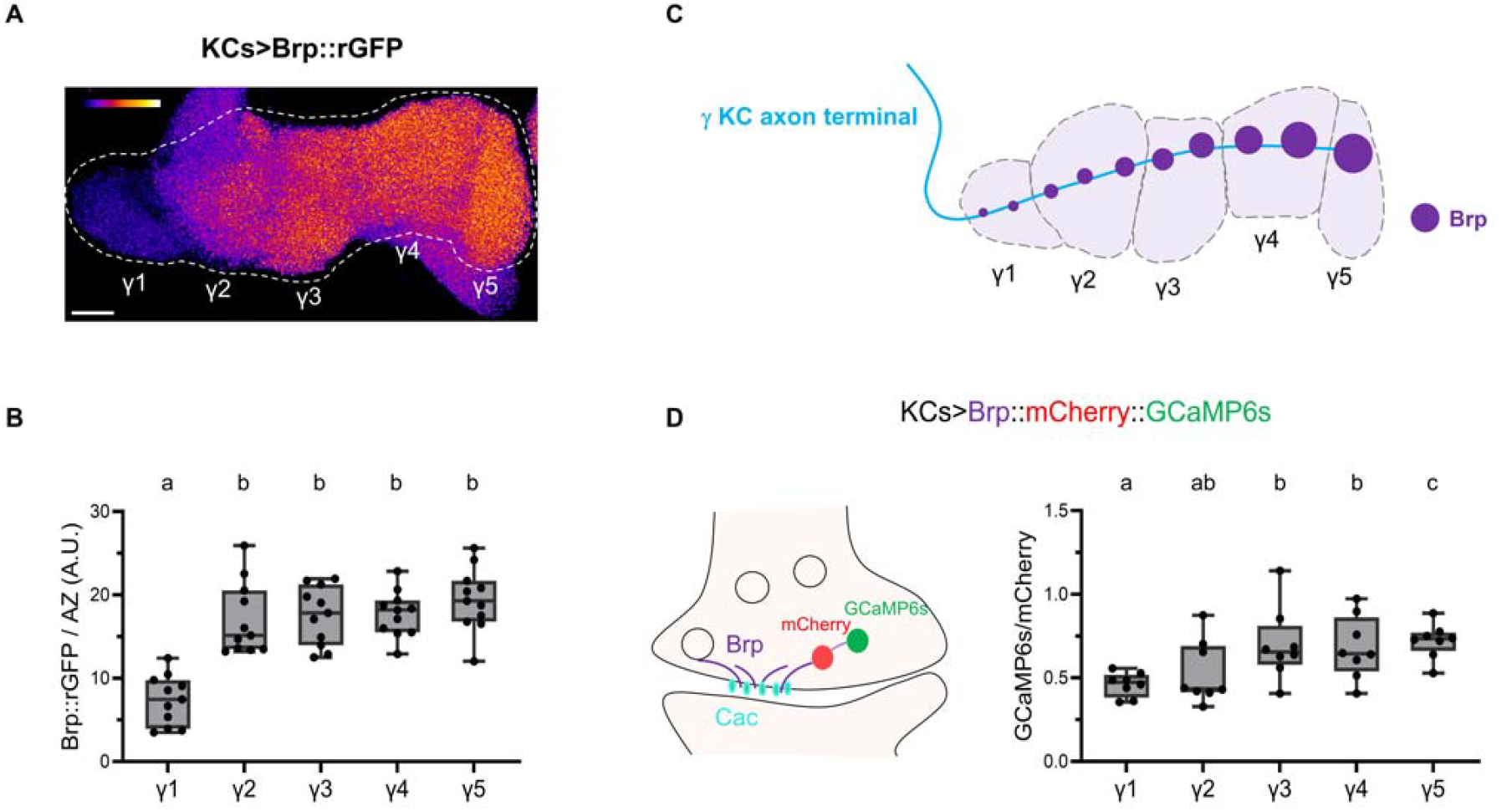
Intracellular active zone heterogeneity among MB compartments. (**A**) Brp::rGFP visualized in KCs. The horizontal lobe of the MB was imaged. Brp::rGFP was visualized using R13F02-GAL4.(**B**) Median intensity of Brp::rGFP clusters in each γ lobe compartment. R13F02-GAL4 was used. Kruskal-Wallis test. n = 11. γ1 vs. γ2: P = 0.0012; γ1 vs. γ3: P = 0.0004; γ1 vs. γ4: P = 0.0004; γ1 vs. γ5: P < 0.0001. (**C**) Diagram showing compartments in the MB γ lobe and the intracellular Brp localization in a single γ KC. (**D**) Basal Ca^2+^ concentration near the active zones varies by compartments. The left panel shows the schematic of the Brp::mCherry::GCaMP6s ratio-matric sensor. GCaMP6s sensor is fused to mCherry and Brp^short^. The sensor was expressed by R13F02-GAL4 and GCaMP6s signal is normalized by mCherry for analysis. Kruskal-Wallis test. n = 8. γ1 vs. γ3: P = 0.0494; γ1 vs. γ4: P = 0.0494; γ1 vs. γ5: P = 0.0155; γ2 vs. γ5: P = 0.0494; Scale bar, 10 μm. Data are represented as box plots showing center (median), whiskers (Min. to Max.). Significant differences (P < 0.05) are indicated by distinct letters or *.

In motor neurons, Brp is well characterized to serve as a scaffold protein accumulating synapse proteins such as voltage-gated Ca^2+^ channels ^17,19,34^. Consistently, we found that the endogenous α subunit of voltage-gated Ca^2+^ channel, Cacophony (Cac) co-localizes with Brp in the γ lobe (Fig. S4, S5). Given the compartmental Brp heterogeneity (Fig. 2B-C), this correlation suggests that the Ca^2+^ concentration is differentially set along the γ KCs. To measure the basal Ca^2+^ concentration nearby AZs, we expressed Brp^short^::mCherry::GCaMP6s, a ratiometric Ca^2+^ sensor fused to mCherry and truncated Brp ^35^, in KCs. We live-imaged the MB and found that basal GCaMP signals were compartmentally distinct and gradually increased towards the γ5 compartment (Fig. 2D). This compartmental pattern of GCaMP signals nicely mirrored the Brp heterogeneity. Taken together the colocalization of Brp and Cac, these results suggest that the Brp accumulation per AZ is set compartmentally, differentiating the basal Ca^2+^ influx via Ca^2+^ channels at AZs ^36^.

### The state-dependent regulation of Brp compartmental heterogeneity

As different MB compartments are functionally coordinated and integrate internal states such as nutritional states, sleep need and aging ^10,37–41^, we examined whether KCs adapt the synaptic structures upon physiological changes. We calculated the variance of the compartments’ log-transformed medians to measure the heterogeneity level. Therefore, the absolute Brp::rGFP intensity does not affect the variance (Fig. 3A). Strikingly, the compartmental heterogeneity in γ KCs significantly decreased upon food deprivation for 48 hours and recovered after refeeding (Fig. 3B, Fig. S8A). In contrast, we did not find a significant effect by aging for 30 d (Fig. S6). These results suggest that acute stress can drive local adjustment of AZ structures within KCs, reflecting reorganized compartmental activities.

**Figure 3.**
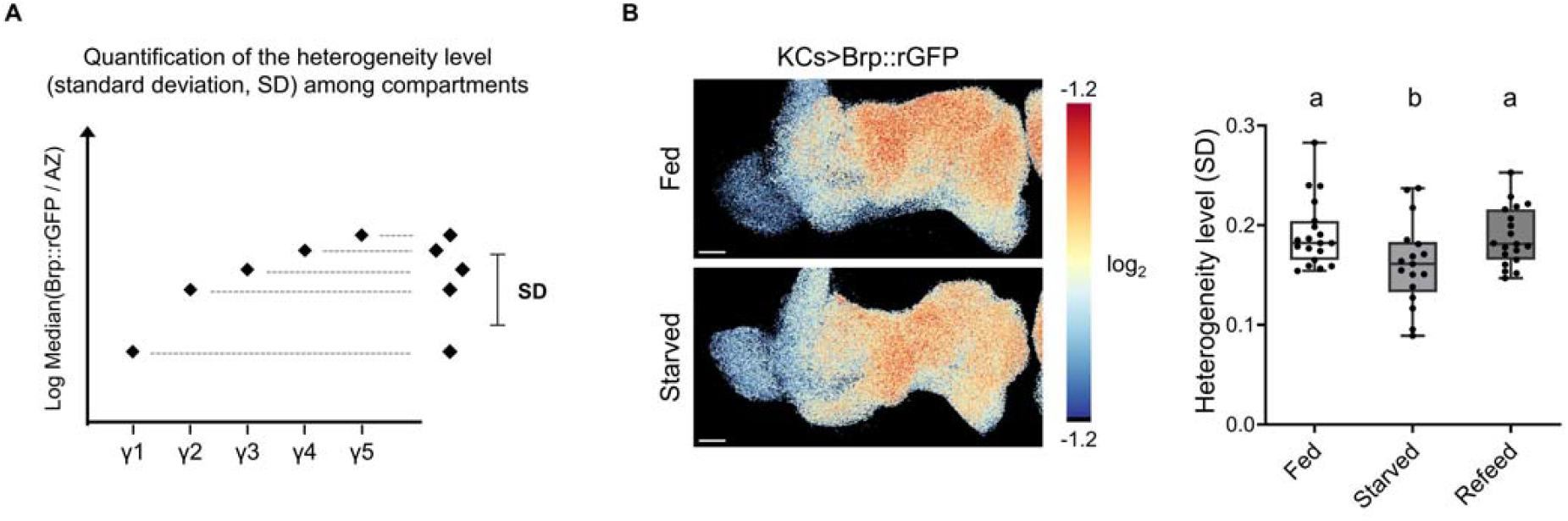
Acute starvation decreases the Brp::rGFP heterogeneity level among γ compartments. (**A**) Standard deviation (SD) as an indicative of Brp::rGFP heterogeneity level among compartments. The medians of Brp::rGFP intensity of clusters in each compartment were log-transformed to calculate the relative ratio of intensity among compartments. SD is calculated from five log-transformed medians for each brain sample. (**B**) Starvation for 48 hours reduced the Brp heterogeneity level in KCs. Pseudo color in the images represent the value of log_2_(pixel intensity/mean pixel intensity in the γ lobe). Pseudo color range: -1.2 to 1.2. The heterogeneity level (SD) was quantified for individual brain samples. Ordinary one-way ANOVA. Fed (n = 19) vs. Starved (n = 17): P = 0.0400; Starved (n = 17) vs. Refeed (n = 19): P = 0.0400; Fed (n = 19) vs. Refeed (n = 19): P = 0.7838; Scale bar, 10 μm. Data are represented as box plots showing center (median), whiskers (Min. to Max.). Significant differences (P < 0.05) are indicated by distinct letters or *.

### Octopamine input underlies the Brp compartmental heterogeneity

Each compartment receives dopamine projections from structurally and functionally distinct subsets of dopaminergic neurons (Fig. S7A) ^9^. We therefore examined whether dopamine inputs determine the compartmental Brp accumulation by knocking-down dopamine receptors (DopR1, DopR2 and D2R) in KCs using RNA interference (RNAi) ^42,43^. None of these disruptions showed a significant effect on the compartmental heterogeneity (Fig. S7B, Fig. S8B).

The octopaminergic neurons (OANs), particularly OA-VPM3 and OA-VPM4 neurons, are known to project to the MB and concentrate their innervations in the γ1 compartment ^9,44^. We characterized presynaptic components of OA-VPM3/4 visualized by Brp::rGFP and nSyb::CLIP (neuronal synaptobrevin, nSyb; synaptic vesicle marker) and found they were densely localized to γ1 compared to the other compartments, complementary to the Brp compartmental heterogeneity of KCs (Fig. 4A-4D). These results suggest localized synaptic output of OANs along the γ lobe, potentially underlying the Brp compartmental heterogeneity.

**Figure 4.**
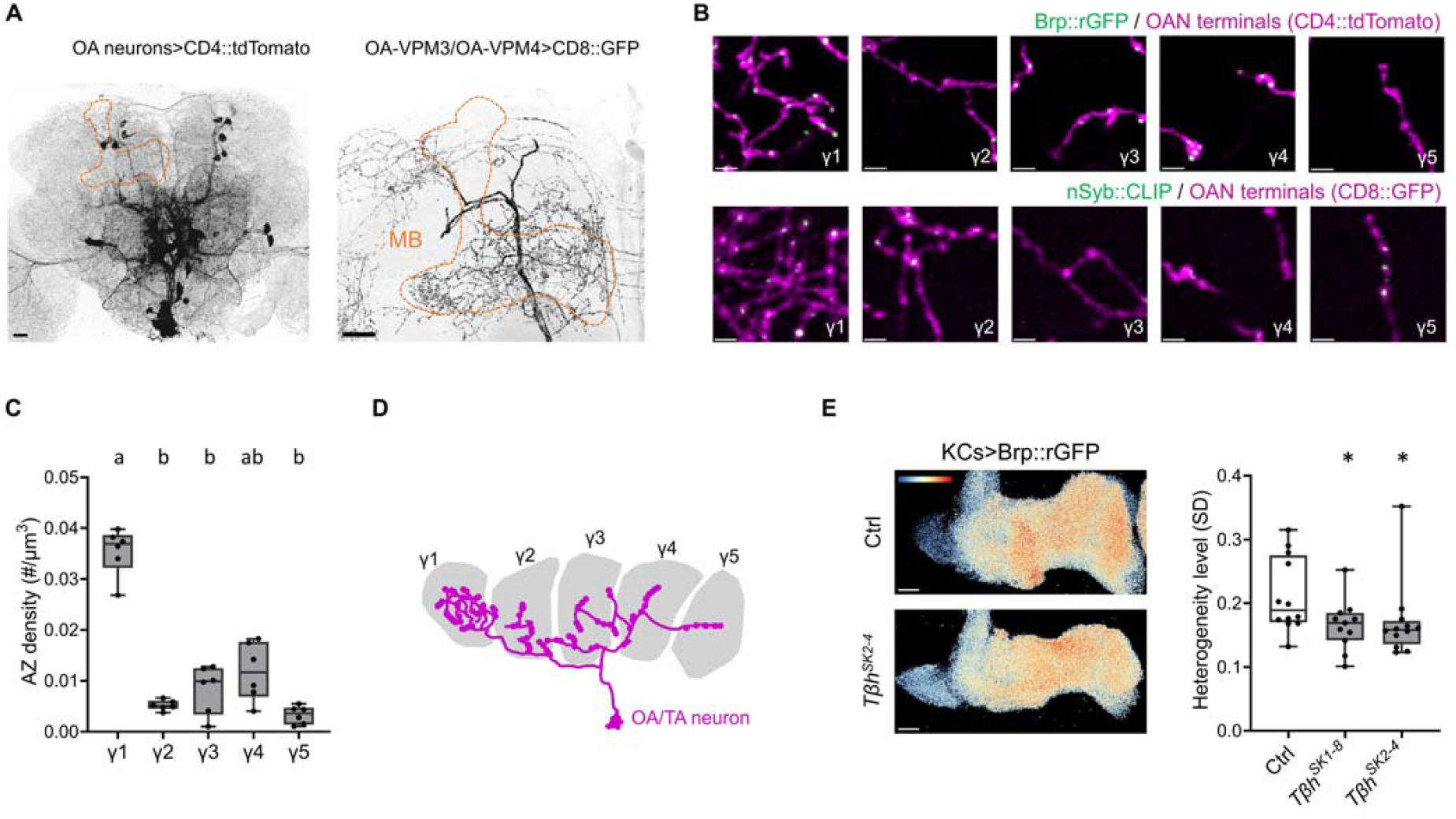
Octopamine controls the AZ structure in KCs. (**A**) OANs in the adult brain. OANs were labeled by Tdc2-GAL4 driven CD4::tdTomato in the left panel. OA-VPM3/4 neurons were labeled with MB022B-GAL4 ^9^ driven CD8::GFP (right panel). Dashed lines indicate the MB. Scale bars, 20 µm. (**B**) AZ number and synaptic vesicle localization of OANs along the γ lobe. GFP_1-10_ and CD4::tdTomato were expressed using Tdc2-GAL4. nSyb::CLIP labeling synaptic vesicles and CD8::GFP were expressed using MB022B-GAL4. Scale bars, 2 µm. Images were cropped from the same MB and are identical in size. (**C**) AZ density of OANs across γ compartment. Brp::rGFP was visualized using Tdc2-GAL4, and cluster density was quantified. n = 6 brains. Kruskal-Wallis test. γ1 vs. γ2: P = 0.0066; γ1 vs. γ3: P =0.0319; γ1 vs. γ5: P =0.0004. (**D**) Schematic showing the innervation pattern of OANs along the γ lobe. (**E**) Reduced Brp compartmental heterogeneity level in *Tβh* mutants. Representative images of Brp::rGFP in control and *Tβh*^*SK2-4*^ background are shown. Pseudo color in the images represent the value of log2(pixel intensity/mean pixel intensity in the γ lobe). Pseudo color range: -1.0 to 1.0. Scale bars, 10 µm. Kruskal-Wallis test. Ctrl (n = 12) vs. *Tβh*^*SK1-8*^ (n = 11) and *Tβh*^*SK2-4*^ (n = 12). Ctrl vs. *Tβh*^*SK1-8*^: P = 0.0442; Ctrl vs. *Tβh*^*SK2-4*^: P = 0.0228. *P < 0.05. Data are represented as box plots showing center (median), whiskers (Min. to Max.). Significant differences (P < 0.05) are indicated by distinct letters or *. Kruskal-Wallis test.

To examine the contribution of octopamine to the Brp accumulation within KCs, we measured the Brp compartmental heterogeneity in the mutants of tyramine β-hydroxylase (*Tβh*), the enzyme that catalyzes the synthesis of octopamine from tyramine ^45^. We generated two new *Tβh* null alleles: insertion mutant *Tβh*^*SK1-8*^ and deletion mutant *Tβh*^*SK2-4*^ using CRISPR/Cas9-mediated mutagenesis. We visualized Brp::rGFP in KCs of these mutants and found the significantly decreased compartmental heterogeneity of the γ KCs (Fig. 4E, Fig. S8C). This suggests that octopamine directly controls the AZ structure in KCs.

To identify the signaling pathway that regulates the Brp compartmental heterogeneity, we knocked-down octopamine/tyramine receptors (including OAMB, Octα2R, Octβ1R, Octβ2R, Octβ3R, OctTyrR, TyrR, TyrRII) in KCs using RNAi ^42,46^. Among all the receptors tested, only Octβ2R knockdown significantly decreased the compartmental heterogeneity (Fig. 5A, Fig. S8D-E). We could not test the effect of Octβ1R and Octβ3R because their deficiency in KCs caused abnormal development of the MB. Octβ2R is coupled to the Gs alpha subunit to stimulate cyclic adenosine monophosphate (cAMP) production ^47^. To test whether cAMP plays a role in Brp accumulation, we knocked-down the adenylate cyclase Rutabaga (Rut) in KCs using RNAi ^42,48–52^. The downregulation of Rut resulted in a significant decrease of the compartmental heterogeneity (Fig. 5B, Fig. S8F), suggesting that octopamine functions through Octβ2R-cAMP pathway.

**Figure 5.**
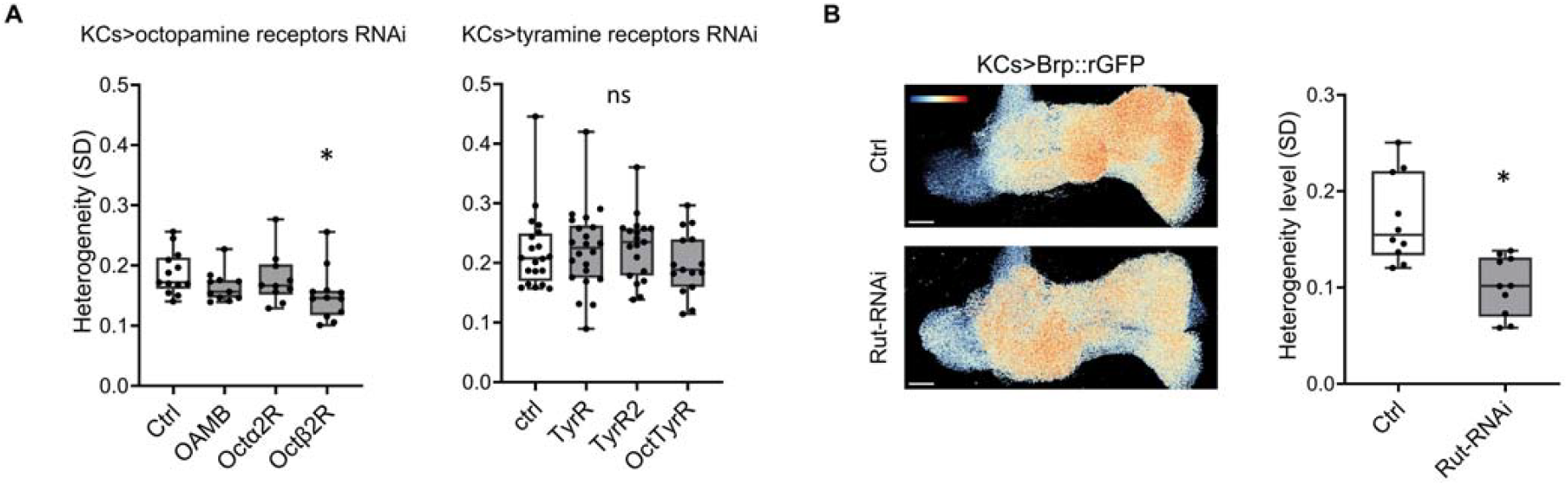
Octβ2R and cAMP underly the Brp compartmental heterogeneity. (**A**) Knockdown of Octβ2R reduced the Brp heterogeneity level in γ KCs. Receptors are knocked-down in γ KCs specifically using RNAi with R13F02-GAL4. Kruskal-Wallis test. Ctrl (n = 13) vs. OAMB (n = 11), Octα2R (n = 10) and Octβ2R (n = 12): P= 0.0228 for Ctrl vs. Octβ2R. Ctrl (n = 20) vs. TyrR (n = 22), TyrR2 (n = 22) and OctTyrR (n = 15). *P < 0.05; ns = not significant. (**B**) Knockdown of Rut significantly decreased the Brp heterogeneity level in γ KCs. Representative images showing Brp::rGFP in control and in Rut knockdown group. Pseudo color in the images represent the value of log2(pixel intensity/mean pixel intensity in the γ lobe). Pseudo color range: -1.2 to 1.2. Mann-Whitney U-test. Ctrl (n = 10) vs. Rut-RNAi (n = 10): P = 0.0011. Data are represented as box plots showing center (median), whiskers (Min. to Max.). *P < 0.05 and ns = not significant. Kruskal-Wallis test (A) and Mann-Whitney U-test (B).

### Nutritional states adjust the Brp compartmental heterogeneity through octopamine signaling

The spontaneous firing of octopamine neurons was shown to be decreased upon starvation ^53^. Therefore, we hypothesized state-dependent AZ remodeling through octopamine (Fig. 6A). To test this hypothesis, we starved the *Tβh* mutants for 48 hours and measured the Brp::rGFP heterogeneity in γ KCs. Indeed, there was no significant state-dependent effect on the Brp compartmental heterogeneity in the *Tβh* mutants in contrast to the control experiment (Fig. 6B, Fig. S8G-H). Taken together, we propose that internal states adjust the synaptic output of MB compartments through octopamine signaling that supervise the synaptic structures within KCs.

**Figure 6.**
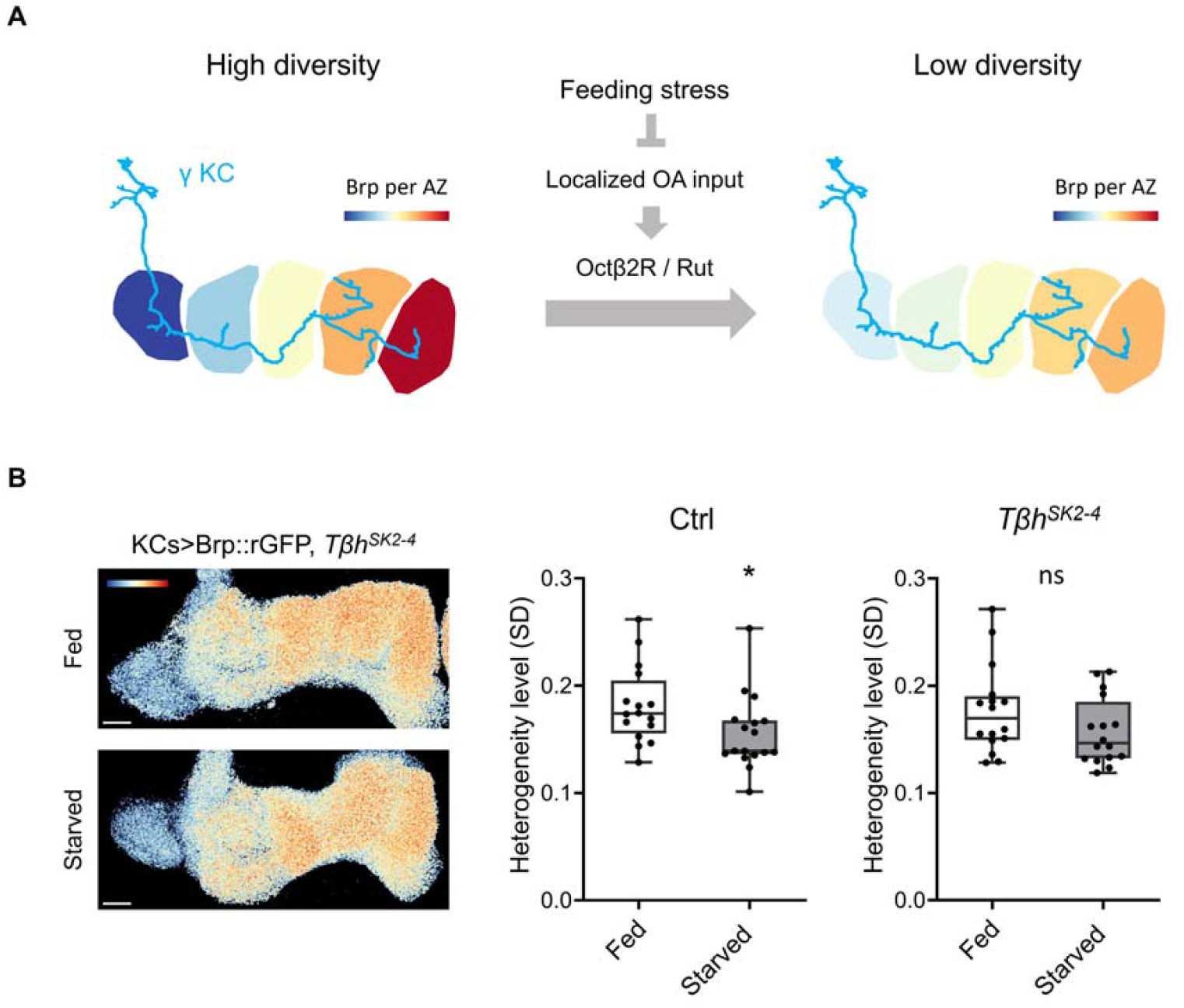
Nutritional states adjust the Brp compartmental heterogeneity through octopamine. (**A**) Graphical summary of the working model. Localized innervation of OANs onto γ KCs regulates the Brp accumulation and forms the Brp compartmental heterogeneity via Octβ2R and cAMP pathway. Nutritional states may modulate OAN activity, thereby adjusting the Brp heterogeneity. (**B**) Starvation for 48 hours did not affect the Brp::rGFP heterogeneity level in *Tβh*^*SK2-4*^ mutant. GFP_1-10_ was expressed in KCs using R13F02-GAL4 in control, or *Tβh*^*SK2-4*^ background flies. Pseudo color in the images represent the value of log2(pixel intensity/mean pixel intensity in the γ lobe). Pseudo color range: -1.2 to 1.2. For the control group, Fed (n = 16) vs. Starved (n = 17): P = 0.0187; For the *Tβh*^*SK2*^ group, Fed (n = 16) vs. starved (n = 17): P > 0.05. Data are represented as box plots showing center (median), whiskers (Min. to Max.). *P < 0.05 and ns = not significant by Mann-Whitney U-test.

## DISCUSSION

By labeling endogenous AZ scaffold protein Brp, we resolved individual AZs at the single-cell level. The substantial intracellular heterogeneity of Brp accumulation within γ KCs is found to be regulated by internal states of the animal. Other than the compartmental pattern of Brp clusters within KCs, our experimental pipeline enables further profiling of other AZ characteristics such as the local AZ density^29^.

How does octopaminergic signaling modify Brp accumulation? This study identified the requirement of Octβ2R and Rut in regulating the AZ structure in KCs (Fig. 5). Octβ2R was shown to stimulate cAMP synthesis ^47^, suggesting that cAMP could play a determining role. Consistent with this idea, *rut* mutant displays abnormal AZ structures ^54^. We propose that the localized innervations of octopamine neurons onto KCs (Fig. 4) results in different cAMP concentrations along the γ lobe. In both KCs and motor neurons, cAMP concentration was shown to be locally regulated ^55–57^. The affinity or stability of Brp to AZs among different compartments is perhaps distinctly set by the cAMP signaling compartmentation^58^. Interestingly, both DopR1 and Octβ2R stimulate cAMP biosynthesis, while the knock-down of DopR1 had no significant effect on the Brp heterogeneity (Fig. S7). This may be due to different downstream targets of the receptors including PKA ^59^.

AZ structures, especially Brp clusters, frequently serve as a proxy to explain synaptic functions ^5–7,17,19,34,60^. Consistent with previous studies, our data suggest that Brp regulates the localization of Ca^2+^ channels in KCs ^17,19,61^. The Brp heterogeneity may result in compartmental basal Ca^2+^ concentrations at AZs (Fig. 2D) and thus modulate spontaneous synaptic vesicle release via the stochastic activity of Ca^2+^ channels ^36^. Indeed, the enrichment of the Ca^2+^ channel is associated to higher spontaneous release frequency in *Drosophila* NMJ ^7,62^. Although the relationship between Brp accumulation and evoked release in KCs seems more complicated since it involves acute and compartmental dopaminergic modulations ^13,63–65^. Unlike evoked activities, spontaneous release of synaptic vesicle persists in the absence of stimulation thereby affecting the states of the animal ^10^.

The Brp compartmental heterogeneity within KCs might directly modulate the MB output through controlling the activity of different MBONs. Each type of MBONs serves as an independent output unit, together regulating a variety of behaviors ^11–14,40,66–68^. Starvation changes the activity balance between MBON-γ1 and MBON-γ5 thus enhances appetitive memory expression ^66,68,69^, and MBON-γ5 and MBON-γ2 oppositely control sleep amount ^70^. Consistently, we found that the compartmental AZ heterogeneity within KCs, measured by Brp, decreased upon starvation (Fig. 3B). Considering the associations between octopamine and feeding behaviors ^53,71–77^, KCs generates behavior adaptations in response to changing internal states and environments by local AZ tuning ^40,78,79^.

## STAR METHOD

### KEY RESOURCES TABLE

| REAGENT or RESOURCE | SOURCE | IDENTIFIER |
| --- | --- | --- |
| **Antibodies** |  |  |
| Anti-Brp | DSHB | Cat#nc82; RRID: AB_2314866 |
| AlexaFluor-568 goat anti-mouse | Invitrogen | Cat#A11004; RRID: AB_2534072 |
| **Chemicals** |  |  |
| SNAP-Surface Alexa Fluor 647 | New England Biolabs | Cat#S9136S |
| CLIP-Surface Alexa Fluor 647 | New England Biolabs | Cat#S9234S |
| OMNIPAQUE 350 | GE Healthcare | Cat#22000AMX02447 |
| Histodenz | Sigma | Cat#D2158 |
| Tetraspeck Microspheres 0.1 mm | Thermo Fisher Scientific | Cat#T7279 |
| NaCl | Nacalai Tesque | Cat#31320-05 |
| KCl | Nacalai Tesque | Cat#28514-75 |
| TES | Nacalai Tesque | Cat#329810-55 |
| Trehalose dihydrate | Tokyo Kasei Kogyo | Cat#T0331 |
| D-glucose | Nacalai Tesque | Cat#16806-25 |
| NaHCO<sub>3</sub> | Nacalai Tesque | Cat#31213-15 |
| NaH<sub>2</sub>PO<sub>4</sub> | Merck KGaA | Cat#106346 |
| CaCl<sub>2</sub> | Sigma-Aldrich | Cat#C5080 |
| MgCl<sub>2</sub> | Sigma-Aldrich | Cat#M2670 |
| **Drosophila strains** |  |  |
| *Brp::SNAP*<sup>26</sup> | Bloomington Drosophila stock center (BDSC) | BDSC 58397 |
| MB008B-GAL4<sup>9</sup> | BDSC | BDSC 68291 |
| MB009B-GAL4<sup>9</sup> | BDSC | BDSC 68292 |
| MB370B-GAL4<sup>9</sup> | BDSC | BDSC 68319 |
| VT64246-GAL4<sup>80</sup> | VDRC | VDRC 204311 |
| GH146-GAL4<sup>81</sup> | BDSC | BDSC 30026 |
| TH-GAL4<sup>82</sup> | BDSC | BDSC 8848 |
| R58E02-GAL4<sup>83</sup> | BDSC | BDSC 41347 |
| R13F02-GAL4<sup>83</sup> | BDSC | BDSC 48571 |
| OK371-GAL4<sup>84</sup> | BDSC | BDSC 26160 |
| UAS-Brp::mCherry::GCaMP6s<sup>35</sup> | BDSC | BDSC 77131 |
| *Cac::sfGFP* | Gratz *et al.*<sup>6</sup> | N/A |
| Tdc2-GAL4<sup>85</sup> | BDSC | BDSC 9313 |
| MB022B-GAL4<sup>9</sup> | BDSC | BDSC 68298 |
| UAS-nSyn::CLIP<sup>26</sup> | BDSC | BDSC 58385 |
| UAS-DopR1-RNAi<sup>52</sup> | BDSC | BDSC 55239 |
| UAS-Dop2R-RNAi<sup>52</sup> | BDSC | BDSC 50621 |
| UAS-DopR2-RNAi<sup>52</sup> | BDSC | BDSC 51423 |
| UAS-OAMB-RNAi<sup>46</sup> | VDRC | VDRC 2861 |
| UAS-Octα2R-RNAi<sup>52</sup> | BDSC | BDSC 50678 |
| UAS-Octβ1R-RNAi<sup>52</sup> | BDSC | BDSC 58179 |
| UAS-Octβ2R-RNAi<sup>52</sup> | BDSC | BDSC 34673 |
| UAS-Octβ3R-RNAi<sup>52</sup> | BDSC | BDSC 62283 |
| UAS-TyrR-RNAi<sup>52</sup> | BDSC | BDSC 25857 |
| UAS-TyrR2-RNAi<sup>52</sup> | BDSC | BDSC 64964 |
| UAS-OctTyrR-RNAi<sup>52</sup> | BDSC | BDSC 28332 |
| UAS-Rut-RNAi<sup>52</sup> | BDSC | BDSC 80468 |
| UAS-CD4::tdTomato | BDSC | BDSC 35841 |
| UAS-CD8::GFP | BDSC | BDSC 32194 |
| *Tβh*<sup>SK1-8</sup> | This study | N/A |
| *Tβh*<sup>SK2-4</sup> | This study | N/A |
| *brp::GFP<sub>11</sub>* | This study | N/A |
| UAS-GFP<sub>1-10</sub> | Kondo et al.<sup>25</sup> | N/A |
| Software |  |  |
| Python | N/A | N/A |
| Graphpad PRISM | Graphpad software | https://www.graphpad.com/ |
| Fiji/Image J | NIH | https://imagej.net/software/fiji/downloads |
| Amira Software | Thermo Fisher Scientific | thermofisher.com/amira-avizo |
| DeconvolutionLab2 | Sage et al.<sup>86</sup> | https://bigwww.epfl.ch/deconvolution/deconvolutionlab2/ |
| CLIJ2 | Haase et al.<sup>87</sup> | https://clij.github.io/ |
| 3D ImageJ Suite | Ollion et al.<sup>88</sup> | https://imagej.net/plugins/3d-imagej-suite/ |

## RESCOURCE AVAILIBILITY

### Lead contact

Further information and requests on resources and reagents should be directed to and will be fulfilled by the lead contact, Dr. Hiromu Tanimoto.

### Material availability

All reagents generated in this study are available upon request from the lead contact.

### Data and code availability

This study did not generate any unique code. All data needed to evaluate the conclusions in the paper are available from the lead contact upon request.

## EXPERIMENTAL MODEL AND SUBJECT DETAILS

### Animal husbandry

Flies were maintained on standard cornmeal food at 25 °C under a 12:12 hours light-dark cycle for all experiments. All flies used for experiments are 3-7 days old adult males. After hatching, adult flies were transferred to fresh food vials and flip every 2-3 days before experiments. In experiments where flies were fasted, the same population of flies was separated before the experiment to ensure the homogeneity. All flies were food-deprived for 48 hours for the experimental group, while provide normal food supply for the control for the same time. The GAL4-UAS system was used to express transgenes. For each transgenic control in RNAi experiments, crossing respective transgenic line with Canton S strain, w1118, or y1w1118 to replace the balancer chromosomes. Flies for wild-type characterization were also crossed to replace balancer chromosomes. Strains used in this study are as indicated in the KEY RESOURCES TABLE.

## METHOD DETAIL

### Transgenic lines and mutants

Two independent null alleles of the *Tβh* gene were obtained by inducing frameshift indels by the transgenic CRISPR/Cas9 system as previously described ^89^. The following 20-bp gRNA sequences were used:

gRNA#1: GATACGTACACCAGTCCGGA

gRNA#2: GGTGAGACGGGACTACCAGC

*Tβh*^*SK1-8*^ was derived from mutagenesis by gRNA#1 and carries the following frameshift insertion: CGTACACCA-----------------------------GgatggacactGGATGGACAG (the inserted sequence is indicated by lower-case characters). *Tβh*^*SK2-4-4*^ was derived from mutagenesis by gRNA#2 and carries the following frameshift deletion: TCCGGATGGA CTGTGAGGTC (the gap is indicated as hyphens). The *brp::GFP*_*11*_ line was generated by targeted insertion of a GFP_11_ cassette into the endogenous *brp* locus just prior to the stop codon as previously described ^25^.

### Sample preparation

Animals used in experiments involving comparisons between control and experimental groups were dissected on the same day. Data were collected from multiple batches of experiments performed on different days. All steps were performed at room temperature otherwise specified. Flies were anesthetized on ice and placed on ice before dissection. The dissection was performed in ice PBS solution to collect the brain without the ventral nerve cord. Brains are subsequently fixed by 2% paraformaldehyde for 1 hour then washed 3×20 min with 0.1% PBT (0.1% Triton X-100 in PBS) in a PCR tube, typically containing 5-6 brains.

For SNAP or CLIP chemical tagging, brains were incubated in SNAP-Surface 647 (1:1000; NEB; S9136S) or CLIP-Surface 647 (1:1000; NEB; S9234S) in 0.3% PBT for 15 min and washed with 3×20 min with 0.1% PBT. Samples are mounted on microscope slides using the appropriate mounting medium (SeeDB2S, SeeDB2G ^90^ or 86% glycerol according to the imaging condition). Brp::SNAP, Brp::rGFP, Cac::sfGFP, CD4::tdTomato and CD8::GFP samples were imaged without immunohistochemistry.

Immunohistochemistry procedures (for anti-Brp staining in Fig. 1B) were carried out as described ^91^. In brief, fixed brains were washed 3×20 min with 0.1% PBT and incubated in 3% normal goat serum (NGS; Sigma-Aldrich; G9023) for 1 hour at room temperature. Antibodies used were as indicated in resource table (Table S1). Brains were incubated in antibody solutions at 4°C for 48 hours for both primary and secondary antibodies. Sub-resolution fluorescent beads (Tetraspeck Microspheres 0.1 mm, Thermo Fisher Scientific, T7279) were imaged for generating experimental point spread function (PSF). The beads solution was diluted with distilled water and sonicated multiple times to eliminate aggregation.

For *in vivo* imaging of PM-Mns in developing pupae, samples were prepared as previously described ^30^. In brief, white prepupal stage pupae (∼ 0 hour APF) were collected and incubated at 25 °C until the experiment. Before imaging, the puparium case was carefully removed with forceps, and the pupa was transferred to an imaging chamber sealed with a cover slip to create an imaging window on the abdomen. For ex vivo imaging of 3^rd^ instar pupal CNS, the CNS was dissected directly from the animal in PBS at room temperature. The CNS sample was then mounted with a spacer between the microscopic glass and cover slip with PBS, The entire process, from dissection to imaging, was performed within 5 minutes.

### Image acquisition

For fixed samples and *ex vivo* imaging, images are acquired using the Olympus FV1200 confocal microscope platform equipped with GaAsP high sensitivity detectors and a 60×/1.42 NA oil immersion objective (PLAPON60XO, Olympus) and a 30×/1.05 NA silicone immersion objective (UPLSAPO30XS, Olympus). For *in vivo* imaging on pupae, images were acquired using the Zeiss LSM 800 series confocal microscopy equipped with a 40×/1.3 NA objective (Plan-Apochromat 40×/1.3 Oil DIC (UV) VIS-IR M27).

For Brp::SNAP chemical tagging, 30×/1.05 NA objective is used with a voxel size of 0.53 µm × 0.53 µm × 0.84 µm (lateral × lateral × axial) to image the whole brain except for the optical lobes. For Brp::rGFP and UAS-CD4::tdTomato samples in all GAL4 types except Tdc2-GAL4, 60×/1.42 NA oil immersion objective was used and 473 nm laser power: 0.1%; 559 nm laser power: 0.1%; scanning speed: 2 µs/pixel with a voxel size of lateral 0.079 × 0.079 × axial 0.370 µm. For Brp::SNAP/Cac::sfGFP images, 60×/1.42 NA objective is used with a voxel size of 0.079 × 0.079 × 0.370 µm. For Tdc2-GAL4 driven UAS4-CD8::GFP and UAS-nSyb::CLIP samples, 30×/1.05 NA objective is used with a voxel size of 0.414 × 0.414 × 0.84 µm. For MB022B-GAL driven UAS-CD8::GFP and UAS-nSyb-CLIP samples, 30×/1.05 NA objective is used with a voxel size of 0.276 × 0.276 × 0.87 µm. For Tdc2-GAL4 driven Brp::rGFP and UAS-CD4::tdTomato, 60×/1.42 NA objective is used with a scanning voxel size of 0.132 × 0.132 × 0.34 µm. For Brp::rGFP *in vivo* imaging of PM-Mns, a 40×/1.3 NA objective is used with a voxel size of 0.0725 × 0.0725 × 0.460 µm. For Brp::rGFP ex vivo imaging, 60×/1.42 NA objective is used with a voxel size of 0.079 × 0.079 × 0.370 µm and 473 nm laser power: 2.0%. For experimental PSF imaging, SeeDB2S immerged beads were scanned in a setting that is 60×/1.42 NA oil immersion objective; 473 nm laser power: 2.0%; 559 nm laser power: 1.5%; voxel size: 0.079 µm × 0.079 µm × 0.370 µm; scanning speed: 4 µs/pixel, to produce multiple bead images.

### *In vivo* calcium imaging

*In vivo* calcium imaging was performed following a previously described protocol ^91^. Briefly, flies were anesthetized on ice for 3 minutes and placed in a custom-made holding dish on a Peltier plate (CP-085, Sinics) maintained at 5°C. The head capsule was fixed to the dish using UV curing optical adhesive (NOA68, Thorlabs). To minimize brain movement, the proboscis was glued to the capsule A small window was opened on the top of the head capsule, and the exposed area was filled with *Drosophila* saline solution (final concentration: 103 mM NaCl, 3 mM KCl, 5 mM TES, 8 mM Trehalose dihydrate, 10 mM D-glucose, 26 mM NaHCO_3_, 1 mM NaH_2_PO_4_, 1.5 mM CaCl_2_, 4 mM MgCl_2_, adjust to pH ∼7.2). Air sacs and fat bodies covering the brain surface were carefully removed. Live imaging was conducted using a laser scanning confocal microscope equipped with GaAsP detectors (A1R, Nikon) and a 30×/1.1 NA water immersion objective (Apo LWD 25×, Nikon). GCaMP6s and mCherry were sequentially excited at 488 and 561 nm, respectively. Emission light was collected using dichroic mirrors and emission filters (BP500–550 and BP570-620) onto GaAsP detectors. The MB γ lobe was scanned at a pixel size of 0.5μm/pixel (512 × 128 pixels) at a frame rate of 1 Hz using line scans with 16× averages in the resonant scanning mode. The pinhole was set to 2.5 AU (561 nm). For recording, images were sequentially acquired over 40 s and saved for later image processing.

### Data processing and analysis

PSF images were processed in the Amira software (Thermo Fisher Scientific). Using the Extract Point Spread Function module, PSFs extracted from each image were averaged into a single PSF, which was later used for image deconvolution (resized voxel size: 0.079 µm × 0.079 µm × 0.370 µm; image size: 32 pixels × 32 pixels × 21 slices). Image deconvolution was performed using the Richardson-Lucy iterative non-blind algorithm in the Fiji plugin DeconvolutionLab2 ^86^, or with CLIJ2 GPU-based Richardson-Lucy deconvolution ^87^.

AZ detection was performed as described ^29^ in a compartmental manner. In brief, a sub-image stacks were cropped from every MB compartment in the original image. Compartments were manually identified referencing the CD4::tdTomato membrane signal. An intensity threshold was applied to reduce background noise. Sub-stacks were processed by the 3D maxima finder and the 3D spot segmentation function in the 3D suite plugin ^88^ in Fiji. 3D maxima were detected for each Brp::rGFP cluster and used as start points for pixel clustering using the 3D spot segmentation function. This process created 3D ROIs enclosing individual Brp clusters. ROIs were then used to extract Brp::rGFP signal intensities independently for each cluster. Detection precision was optimized by comparing detection results with manually defined ground truths as previously described ^29^.

For calcium imaging, ROIs were manually drawn for each compartment in time-series image stacks. GCaMP6s signal was normalized by mCherry in each time-frame using the image calculator in Fiji. The mean intensity was calculated in each compartment by averaging 7 frames without any odor stimulation as the basal level.

## QUANTIFICATION AND STATISTICAL ANALYSIS

### Statistical analysis

Statistical analyses were performed using GraphPad Prism version 8, 9 and 10. Summarized data are represented as box plots showing center (median), whiskers (Min. to Max.). For comparison between more than two groups, Kruskal-Wallis test with original False Discovery Rate (FDR) method of Benjamini and Hochberg correction was applied. Desired false discovery rate was set to 0.05. For comparison between two groups, unpaired nonparametric two-sided Mann-Whitney U-test was used. Pearson’s correlation coefficient (R) was calculated in Python.

## ACKNOWLEDGMENTS

We thank all lab members of Tanimoto Lab at Tohoku University for valuable discussions. We thank Dr. Kate O’Connor-Giles (Brown University), Vienna Drosophila Resource Center (VDRC) and Bloomington Drosophila Stock Center (BDRC) for transgenic flies, Ayano Wu (Tohoku University) for graphical design.

## FUNDING

Ministry of Education, Culture, Sports, Science and Technology 22H05481 (HT)

Ministry of Education, Culture, Sports, Science and Technology 22KK0106 (HT)

Ministry of Education, Culture, Sports, Science and Technology 20H00519 (HT)

Japan Science and Technology Agency JPMJSP2114 (HW)

Tohoku University Research Program ‘Frontier Research in Duo’ (HT)

## AUTHOR CONTRIBUTIONS

Conceptualization, HW, NY and HT

Investigation, HW, SE, KS, AA, KJ

Methodology, HW, NY, SE, SK and YM

Writing – Original Draft, HW, SE and HT

Writing – Review & Editing, all authors

Funding Acquisition, HW and HT

Resources, KJ and SK

Supervision, DW, NY, SK and HT

## COMPETING INTERESTS

Authors declare that they have no competing interests.

## SUPPLEMENTARY INFROAMTION

**Figure S1.**
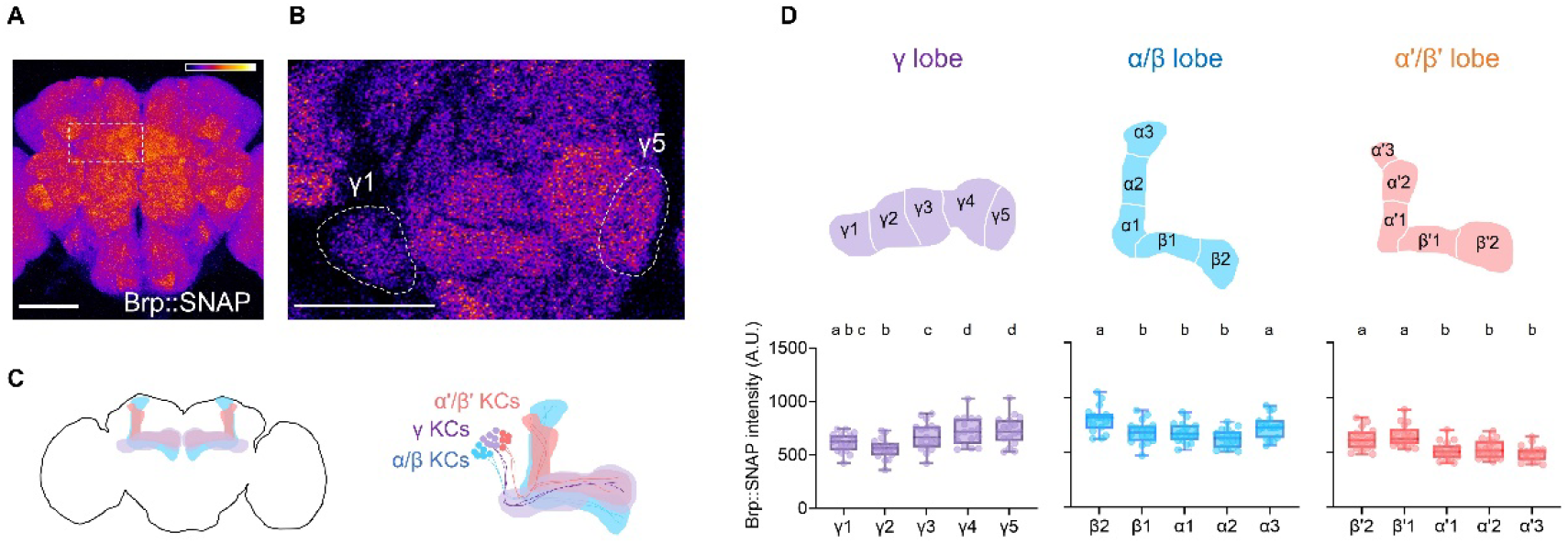
Heterogeneous Brp enrichment in the MB. (**A**) SNAP chemical tagging labeled endogenous Brp in the brain. Scale bar, 100 μm. The dash line area in indicates the zoomed-in area showed in (B). (**B**) Intensity difference of Brp::SNAP between γ1 and γ5 compartment on the same imaging plate. Scale bar, 50 μm. (**C**) Schematic of the MB. The MB comprise three lobes based on the projection patterns of three KC subtypes: γ KCs, α’/β’ KCs and α/β KCs. (**D**) Brp::SNAP intensity difference across compartments. Schematic drawings above indicate compartments of each lobe. Data are represented as box plots showing center (median), whiskers (Min. to Max.). Significant differences (P < 0.05) are indicated by distinct letters. Kruskal-Wallis test.

**Figure S2.**
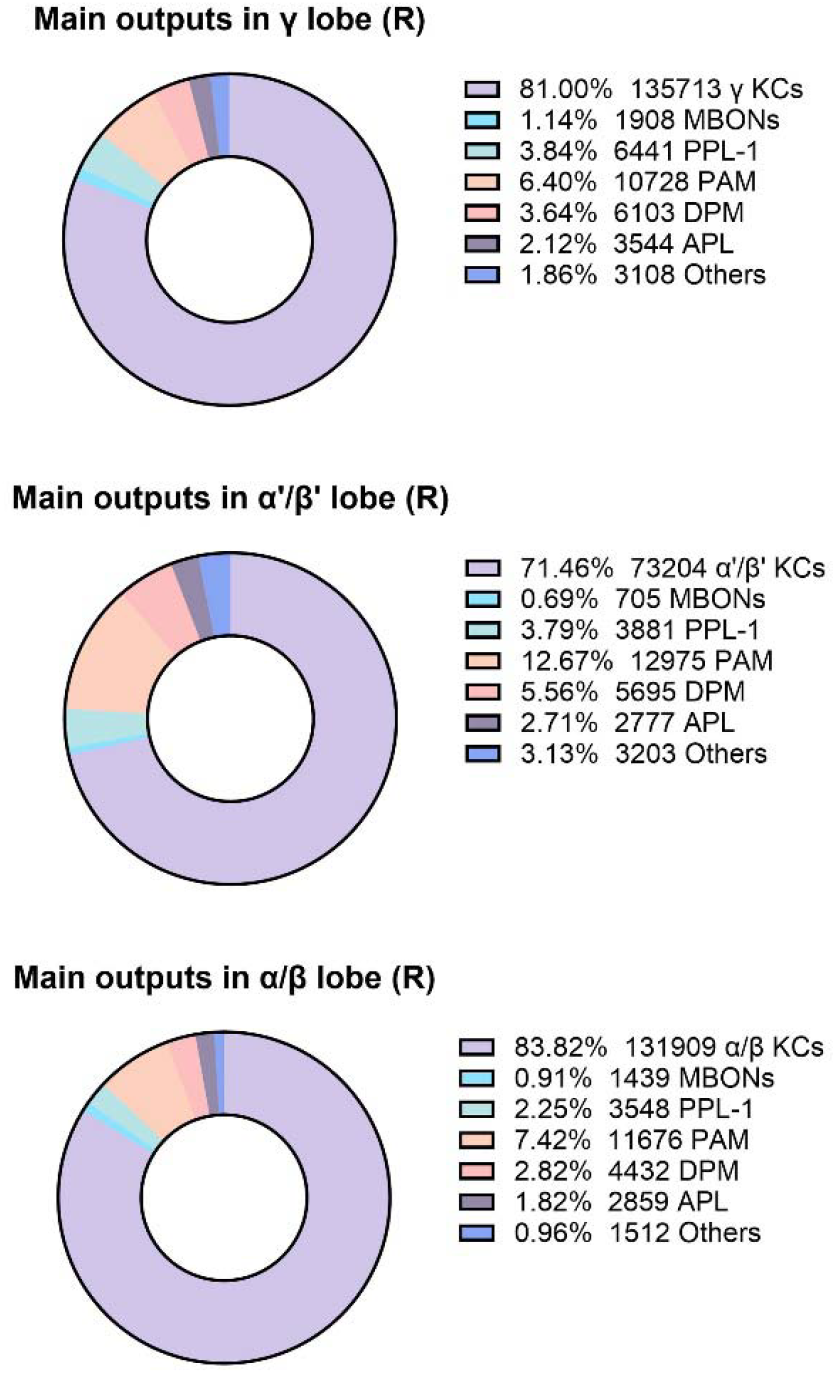
Pre-synapses in the MB are predominantly from KCs. Pie charts showing the composition of pre-synapses in each of the three MB lobes. Data is from the hemibrain online data but not all the cell types are listed. The number of pre-synapse and the percentage are indicated for each cell type, including KCs, MBONs PPL-1 neurons, PAM neurons, dorsal paired medial (DPM) neuron and anterior paired lateral (APL) neuron.

**Figure S3.**
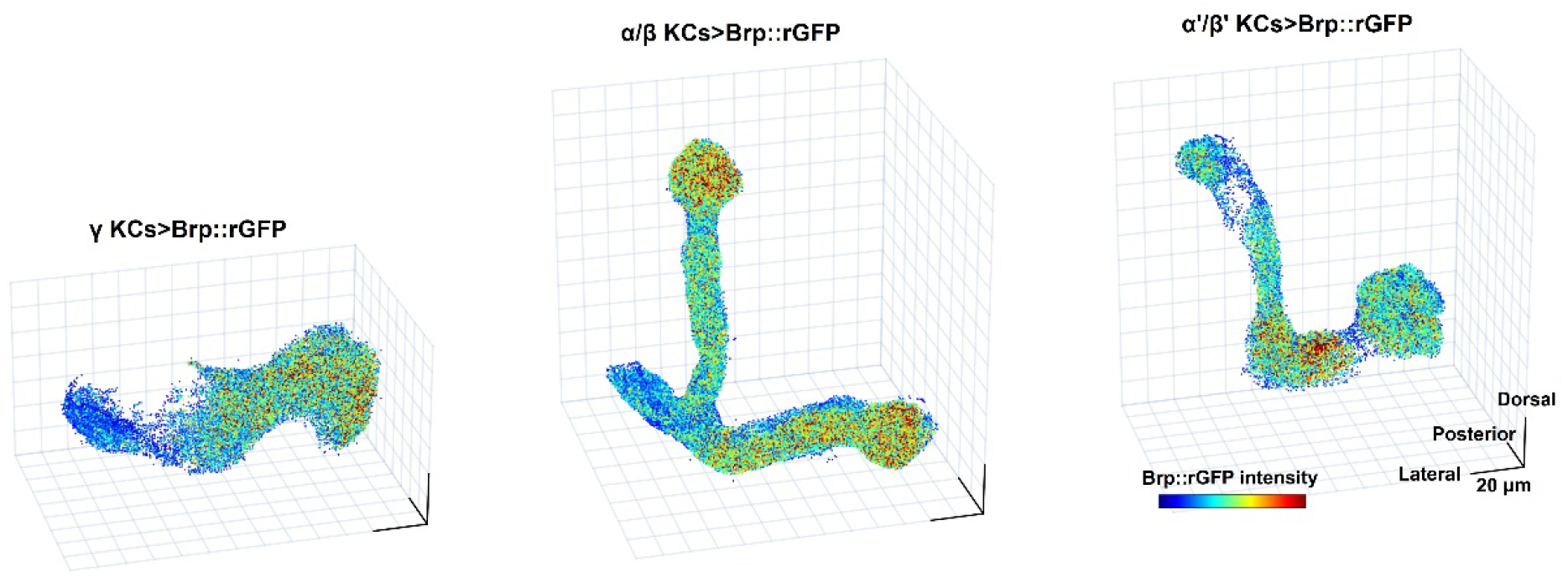
Brp compartmental heterogeneity in different KC subtypes. The 3D reconstruction of Brp::rGFP clusters, colored by Brp::rGFP intensity. Brp::rGFP is visualized in γ KCs using MB009B-GAL4, in α/β KCs using MB008B-GAL4 and in α’/β’ KCs using MB0370B-GAL4. Scale bars, 20 μm.

**Figure S4.**
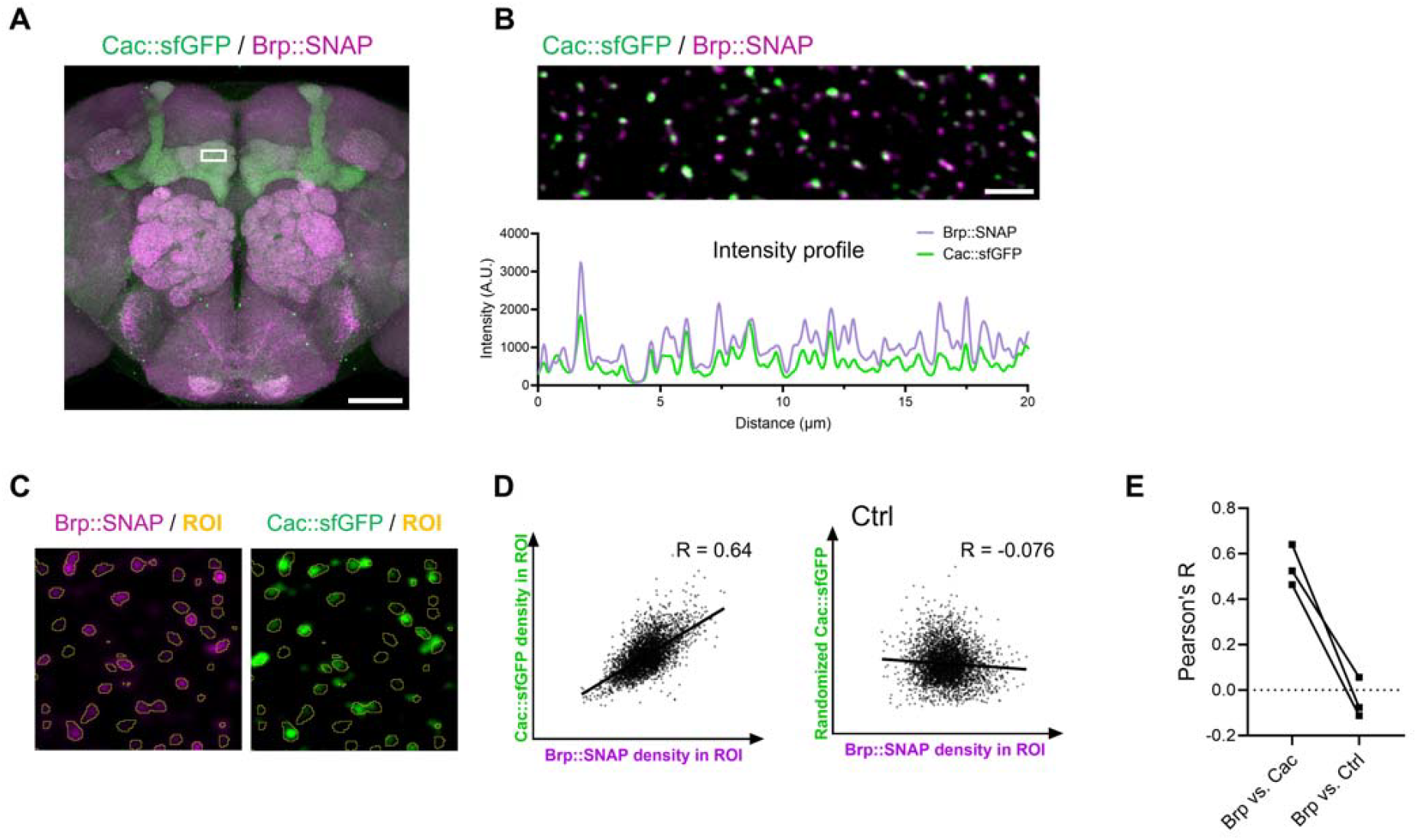
Brp is associated with Ca^2+^ channels in the fly brain. (**A**) Co-labeling of Cac::sfGFP (green) and Brp::SNAP (magenta) in an adult brain. The write box indicates the zoomed-in areas shown in (B). Scale bar, 50 μm. (**B**) Co-localization of Brp::SNAP and Cac::sfGFP signals. Signal intensity profiles of both Brp::SNAP (green) and Cac::sfGFP (magenta) in the image were plotted below. Scale bar, 10 μm. (**C**) Correlation between Brp::SNAP (green) and Cac::sfGFP (magenta) signal intensities. The image is a selected area from γ5. Yellow circles show the 3D ROIs generated by segmenting Brp::SNAP signals. A loose setting was applied to include surrounding pixels. The same ROI set was used to quantify the signal density (total grey value divided by the ROI volume) for both Brp::SNAP and Cac::sfGFP. (**D**) Scatter plot showing the correlation between Brp::SNAP and Cac::sfGFP signal intensities in an image sample. A 180° rotated Cac::sfGFP image was used as a control (see also Fig. S4). Pearson’s correlation coefficient (R) is shown. (**E**) Pearson correlation coefficient (R) from three individual γ lobes showing the correlation between Brp::SNAP and Cac::sfGFP signal intensities. Data are represented as box plots showing center (median), whiskers (Min. to Max.). Significant differences (P < 0.05) are indicated by distinct letters. Kruskal-Wallis test.

**Figure S5.**
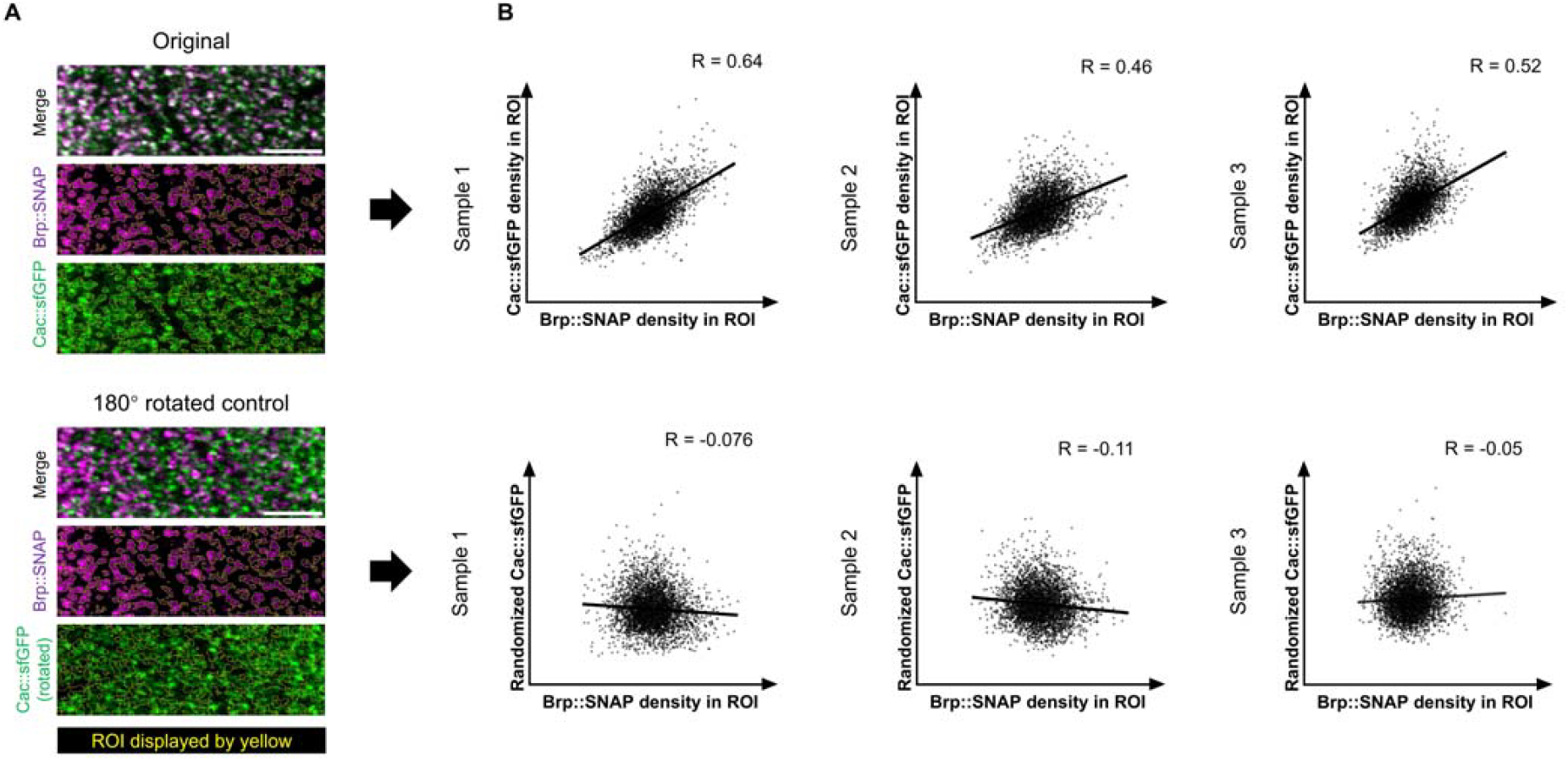
Correlation between Brp::SNAP and Cac::sfGFP signals intensities. (**A**) Correlation analysis of Brp::SNAP and Cac::sfGFP signal intensities. ROIs are generated using 3D spot segmentation method without a watershed process. Relatively loose setting was applied on Brp::SNAP images to generate wide ROIs. The same ROI set was used to calculated signal intensities in both Brp::SNAP and Cac::sfGFP channels. The 180° rotated version of Cac::sfGFP image was used as the control. A total of 4000 ROIs were analyzed, and Pearson correlation coefficient (R) values were calculated for each sample. Scale bars, 5 μm. (**B**) Scatter plots showing the correlation between Brp::SNAP and Cac::sfGFP signal intensities (left) and control (right) in different brain samples. R value is indicated for each sample.

**Figure S6.**
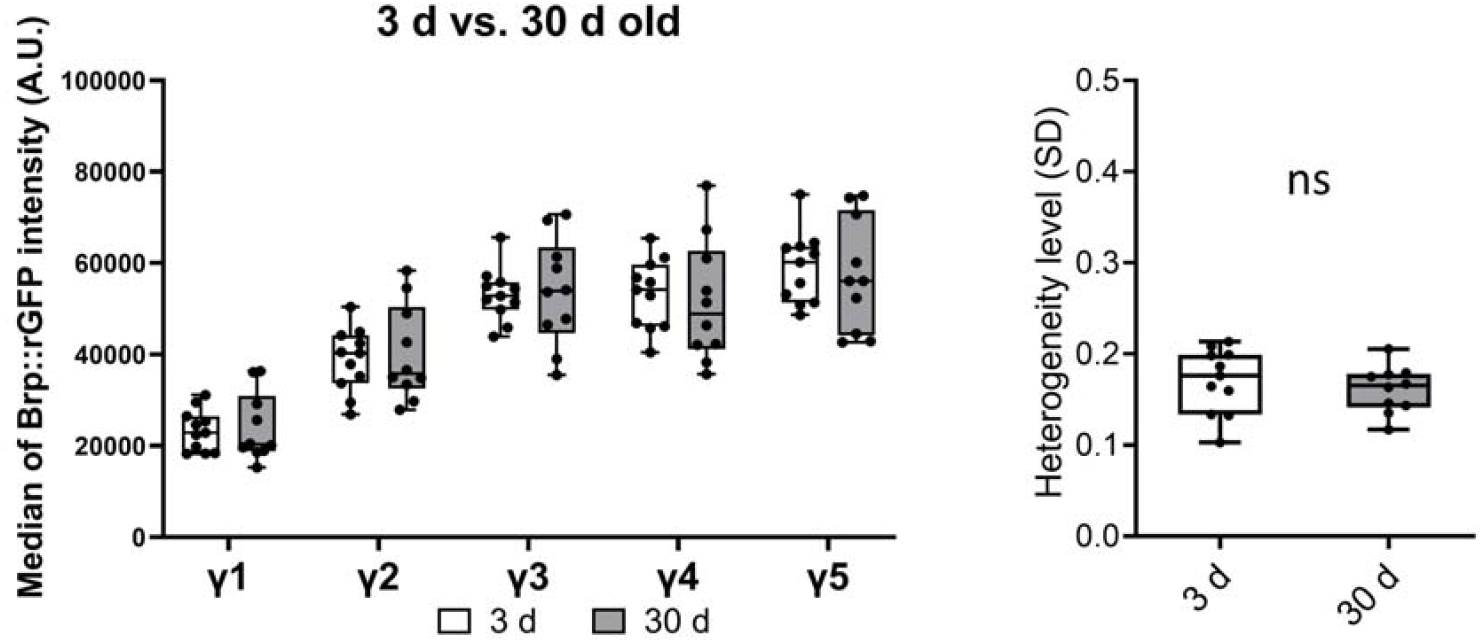
Brp compartmental heterogeneity in aging flies. Median intensity of Brp::rGFP clusters and the Brp::rGFP heterogeneity level were compared between 3 days old and 30 days old flies. Mann-Whitney U-test, 3 d (n = 11) vs. 30 d (n = 10): p > 0.05. Data are represented as box plots showing center (median), whiskers (Min. to Max.). ns = not significant by Mann-Whitney U-test.

**Figure S7.**
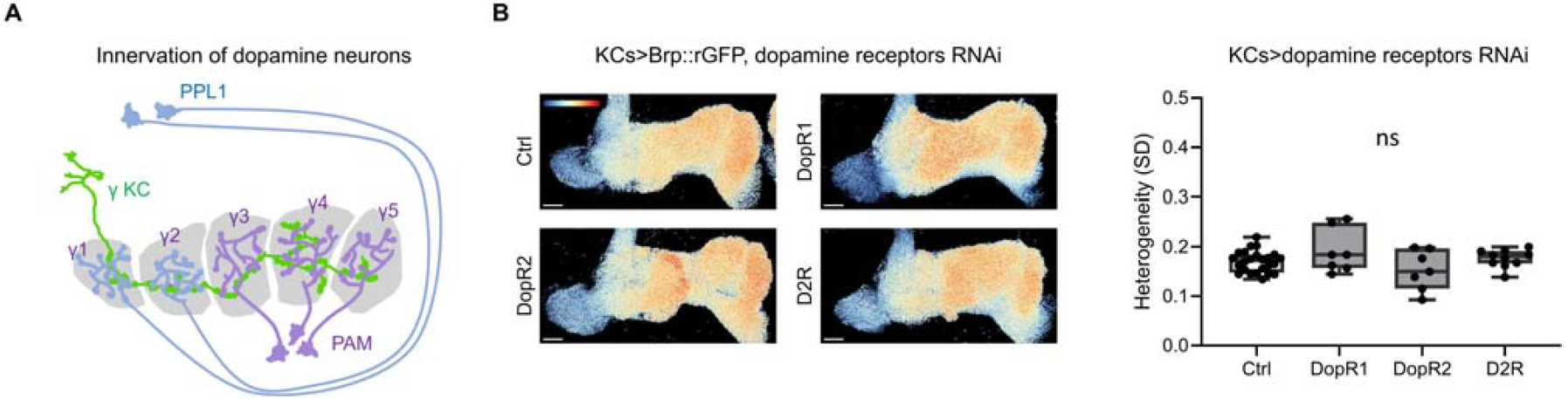
Dopamine receptors knockdown does not affect the Brp compartmental heterogeneity. (**A**) Schematic showing the innervation patterns of dopamine neurons. The γ lobe is innervated by dopamine neurons from the PPL1 and PAM clusters. (**B**) Knockdown of dopamine receptors does not significantly alter the Brp heterogeneity level. Three types of DA receptors, DopR1, DopR2 and D2R were knocked down using RNAi in KCs specifically with R13F02-GAL4. Representative images of Brp::rGFP from each group are shown. Pseudo color in the images represent the value of log2(pixel intensity/mean pixel intensity in the γ lobe). Pseudo color range: -1.3 to 1.3. Scale bar: 20 μm. Ctrl (n = 24) vs. DopR1 (n = 7), DopR2 (n = 7) and D2R (n = 10): P > 0.05. Data are represented as box plots showing center (median), whiskers (Min. to Max.). ns = not significant by Kruskal-Wallis test.

**Figure S8.**
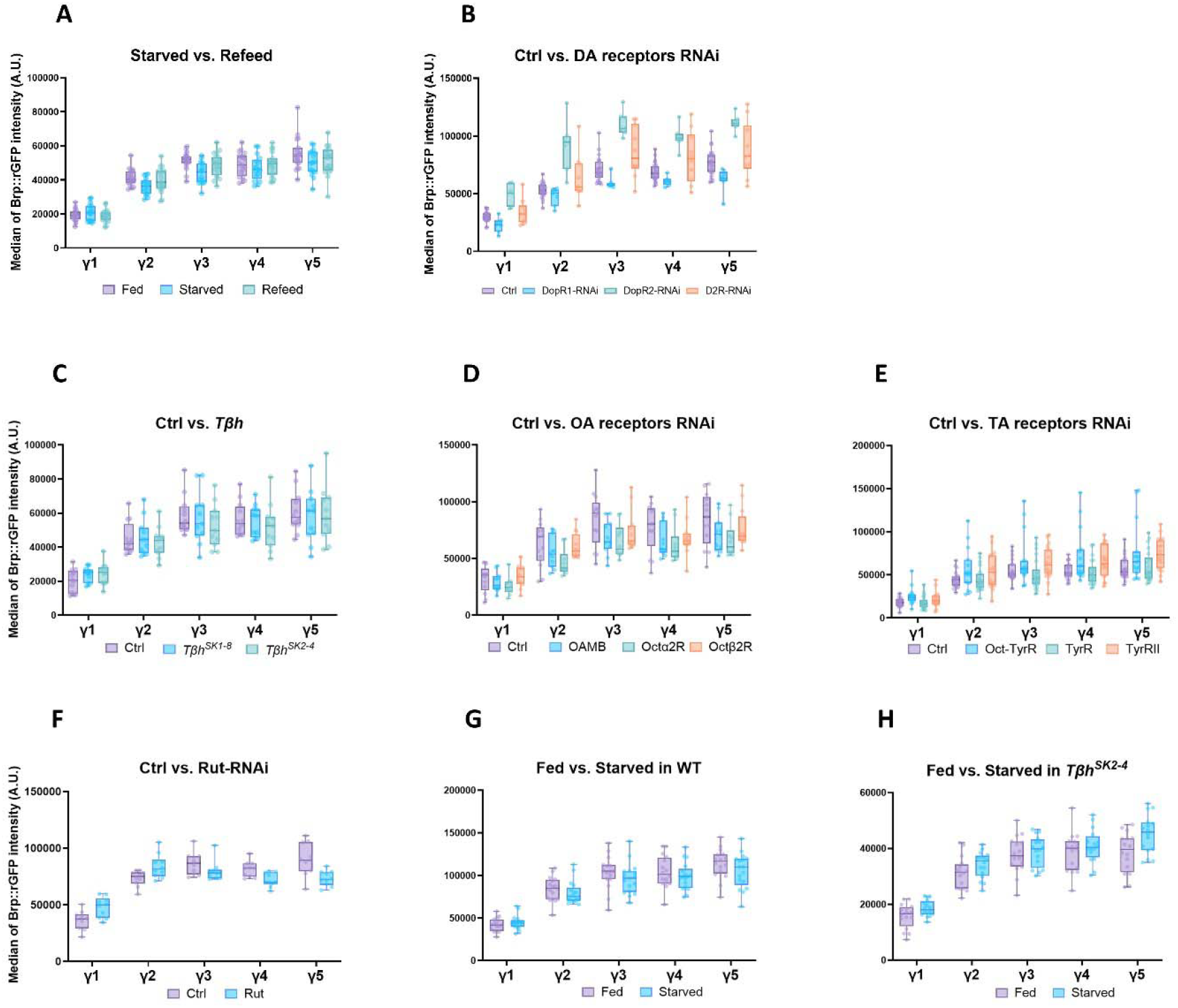
Median intensities of Brp::rGFP clusters in each experiment. (**A**) Starvation and refeeding experiment. (**B**) Dopamine receptor knockdown experiment. (**C**) *Tβh* mutant’s experiment. (**D**) Octopamine receptor knockdown experiment. (**E**) Tyramine receptor knockdown experiment. (**F**) Rut knockdown experiment. (**G**) Starvation experiment. (**H**) Starvation experiment of *Tβh*^*SK2-4*^ mutants. Data are represented as box plots showing center (median), whiskers (Min. to Max.).

## Notes

### Competing Interest Statement

The authors have declared no competing interest.

### Summary of Updates

Modified quantification method explanation in the result as well as the method section. Modified the figure presentation and changed the arrangement.

## REFERENCES

1. O’Rourke, N.A., Weiler, N.C., Micheva, K.D., and Smith, S.J. (2012). Deep molecular diversity of mammalian synapses: why it matters and how to measure it. Nat Rev Neurosci 13, 365–379. 10.1038/nrn3170.

2. Cizeron, M., Qiu, Z., Koniaris, B., Gokhale, R., Komiyama, N.H., Fransén, E., and Grant, S.G.N. (2020). A brainwide atlas of synapses across the mouse life span. Science 369, 270–275. 10.1126/science.aba3163.

3. Ehmann, N., Linde S. van de, Alon, A., Ljaschenko, D., Keung, X.Z., Holm, T., Rings, A., DiAntonio, A., Hallermann, S., Ashery, U., et al. (2014). Quantitative super-resolution imaging of Bruchpilot distinguishes active zone states. Nat Commun 5, 4650. 10.1038/ncomms5650.

4. Paul, M.M., Pauli, M., Ehmann, N., Hallermann, S., Sauer, M., Kittel, R.J., and Heckmann, M. (2015). Bruchpilot and Synaptotagmin collaborate to drive rapid glutamate release and active zone differentiation. Front Cell Neurosci 9, 29. 10.3389/fncel.2015.00029.

5. Akbergenova, Y., Cunningham, K.L., Zhang, Y.V., Weiss, S., and Littleton, J.T. (2018). Characterization of developmental and molecular factors underlying release heterogeneity at Drosophila synapses. Elife 7, e38268. 10.7554/elife.38268.

6. Gratz, S.J., Goel, P., Bruckner, J.J., Hernandez, R.X., Khateeb, K., Macleod, G.T., Dickman, D., and O’Connor-Giles, K.M. (2019). Endogenous tagging reveals differential regulation of Ca2+ channels at single AZs during presynaptic homeostatic potentiation and depression. J Neurosci 39, 3068–18. 10.1523/jneurosci.3068-18.2019.

7. Newman, Z.L., Bakshinskaya, D., Schultz, R., Kenny, S.J., Moon, S., Aghi, K., Stanley, C., Marnani, N., Li, R., Bleier, J., et al. (2022). Determinants of synapse diversity revealed by super-resolution quantal transmission and active zone imaging. Nat Commun 13, 229. 10.1038/s41467-021-27815-2.

8. Tanaka, N.K., Tanimoto, H., and Ito, K. (2008). Neuronal assemblies of the Drosophila mushroom body. J. Comp. Neurol. 508, 711–755. 10.1002/cne.21692.

9. Aso, Y., Hattori, D., Yu, Y., Johnston, R.M., Iyer, N.A., Ngo, T.-T., Dionne, H., Abbott, L., Axel, R., Tanimoto, H., et al. (2014). The neuronal architecture of the mushroom body provides a logic for associative learning. Elife 3, e04577. 10.7554/elife.04577.

10. Suárez-Grimalt, R., Kadow, I.C.G., and Scheunemann, L. (2024). An integrative sensor of body states: how the mushroom body modulates behavior depending on physiological context. Learn. Mem. 31, a053918. 10.1101/lm.053918.124.

11. Aso, Y., Sitaraman, D., Ichinose, T., Kaun, K.R., Vogt, K., Belliart-Guérin, G., Plaçais, P.-Y., Robie, A.A., Yamagata, N., Schnaitmann, C., et al. (2014). Mushroom body output neurons encode valence and guide memory-based action selection in Drosophila. eLife 3, e04580. 10.7554/elife.04580.

12. Ichinose, T., Kanno, M., Wu, H., Yamagata, N., Sun, H., Abe, A., and Tanimoto, H. (2021). Mushroom body output differentiates memory processes and distinct memory-guided behaviors. Curr Biol 31, 1294-1302.e4. 10.1016/j.cub.2020.12.032.

13. Cohn, R., Morantte, I., and Ruta, V. (2015). Coordinated and Compartmentalized Neuromodulation Shapes Sensory Processing in Drosophila. Cell 163, 1742–1755. 10.1016/j.cell.2015.11.019.

14. Owald, D., Felsenberg, J., Talbot, C.B., Das, G., Perisse, E., Huetteroth, W., and Waddell, S. (2015). Activity of Defined Mushroom Body Output Neurons Underlies Learned Olfactory Behavior in Drosophila. Neuron 86, 417–427. 10.1016/j.neuron.2015.03.025.

15. Scheffer, L.K., Xu, C.S., Januszewski, M., Lu, Z., Takemura, S., Hayworth, K.J., Huang, G.B., Shinomiya, K., Maitlin-Shepard, J., Berg, S., et al. (2020). A connectome and analysis of the adult Drosophila central brain. Elife 9, e57443. 10.7554/elife.57443.

16. Zheng, Z., Lauritzen, J.S., Perlman, E., Robinson, C.G., Nichols, M., Milkie, D., Torrens, O., Price, J., Fisher, C.B., Sharifi, N., et al. (2018). A Complete Electron Microscopy Volume of the Brain of Adult Drosophila melanogaster. Cell 174, 730-743.e22. 10.1016/j.cell.2018.06.019.

17. Kittel, R.J., Wichmann, C., Rasse, T.M., Fouquet, W., Schmidt, M., Schmid, A., Wagh, D.A., Pawlu, C., Kellner, R.R., Willig, K.I., et al. (2006). Bruchpilot Promotes Active Zone Assembly, Ca^2+^ Channel Clustering, and Vesicle Release. Science 312, 1051–1054. 10.1126/science.1126308.

18. Wagh, D.A., Rasse, T.M., Asan, E., Hofbauer, A., Schwenkert, I., Dürrbeck, H., Buchner, S., Dabauvalle, M.-C., Schmidt, M., Qin, G., et al. (2006). Bruchpilot, a Protein with Homology to ELKS/CAST, Is Required for Structural Integrity and Function of Synaptic Active Zones in Drosophila. Neuron 49, 833–844. 10.1016/j.neuron.2006.02.008.

19. Fouquet, W., Owald, D., Wichmann, C., Mertel, S., Depner, H., Dyba, M., Hallermann, S., Kittel, R.J., Eimer, S., and Sigrist, S.J. (2009). Maturation of active zone assembly by Drosophila Bruchpilot. J Cell Biol 186, 129–145. 10.1083/jcb.200812150.

20. Matkovic, T., Siebert, M., Knoche, E., Depner, H., Mertel, S., Owald, D., Schmidt, M., Thomas, U., Sickmann, A., Kamin, D., et al. (2013). The Bruchpilot cytomatrix determines the size of the readily releasable pool of synaptic vesicles. J Cell Biol 202, 667–683. 10.1083/jcb.201301072.

21. Hallermann, S., Kittel, R.J., Wichmann, C., Weyhersmüller, A., Fouquet, W., Mertel, S., Owald, D., Eimer, S., Depner, H., Schwärzel, M., et al. (2010). Naked Dense Bodies Provoke Depression. J Neurosci 30, 14340–14345. 10.1523/jneurosci.2495-10.2010.

22. Chen, Y., Akin, O., Nern, A., Tsui, C.Y.K., Pecot, M.Y., and Zipursky, S.L. (2014). Cell-type-Specific Labeling of Synapses In Vivo through Synaptic Tagging with Recombination. Neuron 81, 280– 293. 10.1016/j.neuron.2013.12.021.

23. Gärtig, P.-A., Ostrovsky, A., Manhart, L., Giachello, C.N.G., Kovacevic, T., Lustig, H., Chwalla, B., Cachero, S., Baines, R.A., Landgraf, M., et al. (2019). Motor circuit function is stabilized during postembryonic growth by anterograde trans-synaptic Jelly Belly - Anaplastic Lymphoma Kinase signaling. bioRxiv, 841106. 10.1101/841106.

24. Kamiyama, D., Sekine, S., Barsi-Rhyne, B., Hu, J., Chen, B., Gilbert, L.A., Ishikawa, H., Leonetti, M.D., Marshall, W.F., Weissman, J.S., et al. (2016). Versatile protein tagging in cells with split fluorescent protein. Nat. Commun. 7, 11046. 10.1038/ncomms11046.

25. Kondo, S., Takahashi, T., Yamagata, N., Imanishi, Y., Katow, H., Hiramatsu, S., Lynn, K., Abe, A., Kumaraswamy, A., and Tanimoto, H. (2020). Neurochemical Organization of the Drosophila Brain Visualized by Endogenously Tagged Neurotransmitter Receptors. Cell Reports 30, 284-297.e5. 10.1016/j.celrep.2019.12.018.

26. Kohl, J., Ng, J., Cachero, S., Ciabatti, E., Dolan, M.-J., Sutcliffe, B., Tozer, A., Ruehle, S., Krueger, D., Frechter, S., et al. (2014). Ultrafast tissue staining with chemical tags. Proc. Natl. Acad. Sci. 111, E3805–E3814. 10.1073/pnas.1411087111.

27. Jenett, A., Rubin, G.M., Ngo, T.-T.B., Shepherd, D., Murphy, C., Dionne, H., Pfeiffer, B.D., Cavallaro, A., Hall, D., Jeter, J., et al. (2012). A GAL4-Driver Line Resource for Drosophila Neurobiology. Cell Rep. 2, 991–1001. 10.1016/j.celrep.2012.09.011.

28. Waddell, S., Armstrong, J.D., Kitamoto, T., Kaiser, K., and Quinn, W.G. (2000). The amnesiac Gene Product Is Expressed in Two Neurons in the Drosophila Brain that Are Critical for Memory. Cell 103, 805–813. 10.1016/s0092-8674(00)00183-5.

29. Wu, H., Maekawa, Y., Eno, S., Kondo, S., Yamagata, N., and Tanimoto, H. (2024). High-throughput synapse profiling reveals cell-type-specific spatial configurations in the fly brain. bioRxiv, 2024.12.02.626511. 10.1101/2024.12.02.626511.

30. Constance, W.D., Mukherjee, A., Fisher, Y.E., Pop, S., Blanc, E., Toyama, Y., and Williams, D.W. (2018). Neurexin and Neuroligin-based adhesion complexes drive axonal arborisation growth independent of synaptic activity. Elife 7, e31659. 10.7554/elife.31659.

31. McDonald, N.A., Fetter, R.D., and Shen, K. (2020). Assembly of synaptic active zones requires phase separation of scaffold molecules. Nature 588, 454–458. 10.1038/s41586-020-2942-0.

32. Lucy, L.B. (1974). An iterative technique for the rectification of observed distributions. Astronomical J 79, 745. 10.1086/111605.

33. Richardson, W.H. (1972). Bayesian-Based Iterative Method of Image Restoration*. J Opt Soc Am 62, 55. 10.1364/josa.62.000055.

34. Ghelani, T., Escher, M., Thomas, U., Esch, K., Lützkendorf, J., Depner, H., Maglione, M., Parutto, P., Gratz, S., Matkovic-Rachid, T., et al. (2023). Interactive nanocluster compaction of the ELKS scaffold and Cacophony Ca2+ channels drives sustained active zone potentiation. Sci. Adv. 9, eade7804. 10.1126/sciadv.ade7804.

35. Kiragasi, B., Wondolowski, J., Li, Y., and Dickman, D.K. (2017). A Presynaptic Glutamate Receptor Subunit Confers Robustness to Neurotransmission and Homeostatic Potentiation. Cell Reports 19, 2694–2706. 10.1016/j.celrep.2017.06.003.

36. Ermolyuk, Y.S., Alder, F.G., Surges, R., Pavlov, I.Y., Timofeeva, Y., Kullmann, D.M., and Volynski, K.E. (2013). Differential triggering of spontaneous glutamate release by P/Q-, N- and R-type Ca2+ channels. Nat. Neurosci. 16, 1754–1763. 10.1038/nn.3563.

37. Huang, S., Piao, C., Beuschel, C.B., Zhao, Z., and Sigrist, S.J. (2022). A brain-wide form of presynaptic active zone plasticity orchestrates resilience to brain aging in Drosophila. PLOS Biol. 20, e3001730. 10.1371/journal.pbio.3001730.

38. Huang, S., Piao, C., Beuschel, C.B., Götz, T., and Sigrist, S.J. (2020). Presynaptic Active Zone Plasticity Encodes Sleep Need in Drosophila. Curr Biol 30, 1077-1091.e5. 10.1016/j.cub.2020.01.019.

39. Weiss, J.T., Blundell, M.Z., Singh, P., and Donlea, J.M. (2024). Sleep deprivation drives brain-wide changes in cholinergic presynapse abundance in Drosophila melanogaster. Proc. Natl. Acad. Sci. 121, e2312664121. 10.1073/pnas.2312664121.

40. Weiss, J.T., and Donlea, J.M. (2021). Sleep deprivation results in diverse patterns of synaptic scaling across the Drosophila mushroom bodies. Curr. Biol. 31, 3248-3261.e3. 10.1016/j.cub.2021.05.018.

41. Gupta, V.K., Pech, U., Bhukel, A., Fulterer, A., Ender, A., Mauermann, S.F., Andlauer, T.F.M., Antwi-Adjei, E., Beuschel, C., Thriene, K., et al. (2016). Spermidine Suppresses Age-Associated Memory Impairment by Preventing Adverse Increase of Presynaptic Active Zone Size and Release. Plos Biol 14, e1002563. 10.1371/journal.pbio.1002563.

42. Zirin, J., Hu, Y., Liu, L., Yang-Zhou, D., Colbeth, R., Yan, D., Ewen-Campen, B., Tao, R., Vogt, E., VanNest, S., et al. (2020). Large-Scale Transgenic Drosophila Resource Collections for Loss- and Gain-of-Function Studies. Genetics 214, 755–767. 10.1534/genetics.119.302964.

43. Sun, H., Nishioka, T., Hiramatsu, S., Kondo, S., Amano, M., Kaibuchi, K., Ichinose, T., and Tanimoto, H. (2020). Dopamine receptor Dop1R2 stabilizes appetitive olfactory memory through the Raf/MAPK pathway in Drosophila. J Neurosci 40, 1572–19. 10.1523/jneurosci.1572-19.2020.

44. Busch, S., Selcho, M., Ito, K., and Tanimoto, H. (2009). A map of octopaminergic neurons in the Drosophila brain. J Comp Neurol 513, 643–667. 10.1002/cne.21966.

45. Monastirioti, M., Jr., C.E.L., and White, K. (1996). Characterization of Drosophila Tyramine β-HydroxylaseGene and Isolation of Mutant Flies Lacking Octopamine. J Neurosci 16, 3900–3911. 10.1523/jneurosci.16-12-03900.1996.

46. Dietzl, G., Chen, D., Schnorrer, F., Su, K.-C., Barinova, Y., Fellner, M., Gasser, B., Kinsey, K., Oppel, S., Scheiblauer, S., et al. (2007). A genome-wide transgenic RNAi library for conditional gene inactivation in Drosophila. Nature 448, 151–156. 10.1038/nature05954.

47. Maqueira, B., Chatwin, H., and Evans, P.D. (2005). Identification and characterization of a novel family of DrosophilaβLJadrenergicLJlike octopamine GLJprotein coupled receptors. J Neurochem 94, 547–560. 10.1111/j.1471-4159.2005.03251.x.

48. Levin, L.R., Han, P.-L., Hwang, P.M., Feinstein, P.G., Davis, R.L., and Reed, R.R. (1992). The Drosophila learning and memory gene rutabaga encodes a Ca2+calmodulin-responsive adenylyl cyclase. Cell 68, 479–489. 10.1016/0092-8674(92)90185-f.

49. Livingstone, M.S., Sziber, P.P., and Quinn, W.G. (1984). Loss of calcium/calmodulin responsiveness in adenylate cyclase of rutabaga, a Drosophila learning mutant. Cell 37, 205–215. 10.1016/0092-8674(84)90316-7.

50. Han, P.-L., Levin, L.R., Reed, R.R., and Davis, R.L. (1992). Preferential expression of the drosophila rutabaga gene in mushroom bodies, neural centers for learning in insects. Neuron 9, 619–627. 10.1016/0896-6273(92)90026-a.

51. Thum, A.S., Jenett, A., Ito, K., Heisenberg, M., and Tanimoto, H. (2007). Multiple Memory Traces for Olfactory Reward Learning in Drosophila. J Neurosci 27, 11132–11138. 10.1523/jneurosci.2712-07.2007.

52. Perkins, L.A., Holderbaum, L., Tao, R., Hu, Y., Sopko, R., McCall, K., Yang-Zhou, D., Flockhart, I., Binari, R., Shim, H.-S., et al. (2015). The Transgenic RNAi Project at Harvard Medical School: Resources and Validation. Genetics 201, 843–852. 10.1534/genetics.115.180208.

53. LeDue, E.E., Mann, K., Koch, E., Chu, B., Dakin, R., and Gordon, M.D. (2016). Starvation-Induced Depotentiation of Bitter Taste in Drosophila. Curr Biol 26, 2854–2861. 10.1016/j.cub.2016.08.028.

54. Renger, J.J., Ueda, A., Atwood, H.L., Govind, C.K., and Wu, C.-F. (2000). Role of cAMP Cascade in Synaptic Stability and Plasticity: Ultrastructural and Physiological Analyses of Individual Synaptic Boutons in Drosophila Memory Mutants. J Neurosci 20, 3980–3992. 10.1523/jneurosci.20-11-03980.2000.

55. Boto, T., Louis, T., Jindachomthong, K., Jalink, K., and Tomchik, S.M. (2014). Dopaminergic Modulation of cAMP Drives Nonlinear Plasticity across the Drosophila Mushroom Body Lobes. Curr Biol 24, 822–831. 10.1016/j.cub.2014.03.021.

56. Wang, L., Wu, C., Peng, W., Zhou, Z., Zeng, J., Li, X., Yang, Y., Yu, S., Zou, Y., Huang, M., et al. (2022). A high-performance genetically encoded fluorescent indicator for in vivo cAMP imaging. Nat. Commun. 13, 5363. 10.1038/s41467-022-32994-7.

57. Maiellaro, I., Lohse, M.J., Kittel, R.J., and Calebiro, D. (2016). cAMP Signals in Drosophila Motor Neurons Are Confined to Single Synaptic Boutons. Cell Reports 17, 1238–1246. 10.1016/j.celrep.2016.09.090.

58. Zaccolo, M., Zerio, A., and Lobo, M.J. (2021). Subcellular Organization of the cAMP Signaling Pathway. Pharmacol. Rev. 73, 278–309. 10.1124/pharmrev.120.000086.

59. Gervasi, N., Tchénio, P., and Preat, T. (2010). PKA Dynamics in a Drosophila Learning Center: Coincidence Detection by Rutabaga Adenylyl Cyclase and Spatial Regulation by Dunce Phosphodiesterase. Neuron 65, 516–529. 10.1016/j.neuron.2010.01.014.

60. Ehmann, N., Owald, D., and Kittel, R.J. (2017). Drosophila active zones: From molecules to behaviour. Neurosci Res 127, 14–24. 10.1016/j.neures.2017.11.015.

61. Cunningham, K.L., Sauvola, C.W., Tavana, S., and Littleton, J.T. (2022). Regulation of presynaptic Ca2+ channel abundance at active zones through a balance of delivery and turnover. eLife 11, e78648. 10.7554/elife.78648.

62. Williams, C.L., and Smith, S.M. (2018). Calcium dependence of spontaneous neurotransmitter release. J. Neurosci. Res. 96, 335–347. 10.1002/jnr.24116.

63. Handler, A., Graham, T.G.W., Cohn, R., Morantte, I., Siliciano, A.F., Zeng, J., Li, Y., and Ruta, V. (2019). Distinct Dopamine Receptor Pathways Underlie the Temporal Sensitivity of Associative Learning. Cell 178, 60-75.e19. 10.1016/j.cell.2019.05.040.

64. Noyes, N.C., and Davis, R.L. (2023). Innate and learned odor-guided behaviors utilize distinct molecular signaling pathways in a shared dopaminergic circuit. Cell Reports 42, 112026. 10.1016/j.celrep.2023.112026.

65. Abe, T., Yamazaki, D., Hiroi, M., Ueoka, Y., Maeyama, Y., and Tabata, T. (2023). Revisiting the role of cAMP in Drosophila aversive olfactory memory formation. bioRxiv, 2023.06.26.545795. 10.1101/2023.06.26.545795.

66. Tsao, C.-H., Chen, C.-C., Lin, C.-H., Yang, H.-Y., and Lin, S. (2018). Drosophila mushroom bodies integrate hunger and satiety signals to control innate food-seeking behavior. Elife 7, e35264. 10.7554/elife.35264.

67. Hancock, C.E., Rostami, V., Rachad, E.Y., Deimel, S.H., Nawrot, M.P., and Fiala, A. (2022). Visualization of learning-induced synaptic plasticity in output neurons of the Drosophila mushroom body γ-lobe. Sci. Rep. 12, 10421. 10.1038/s41598-022-14413-5.

68. Perisse, E., Owald, D., Barnstedt, O., Talbot, C.B., Huetteroth, W., and Waddell, S. (2016). Aversive Learning and Appetitive Motivation Toggle Feed-Forward Inhibition in the Drosophila Mushroom Body. Neuron 90, 1086–1099. 10.1016/j.neuron.2016.04.034.

69. Pavlowsky, A., Schor, J., Plaçais, P.-Y., and Preat, T. (2018). A GABAergic Feedback Shapes Dopaminergic Input on the Drosophila Mushroom Body to Promote Appetitive Long-Term Memory. Curr Biol 28, 1783-1793.e4. 10.1016/j.cub.2018.04.040.

70. Sitaraman, D., Aso, Y., Jin, X., Chen, N., Felix, M., Rubin, G.M., and Nitabach, M.N. (2015). Propagation of Homeostatic Sleep Signals by Segregated Synaptic Microcircuits of the Drosophila Mushroom Body. Curr. Biol. 25, 2915–2927. 10.1016/j.cub.2015.09.017.

71. Berger, M., Auweiler, K., Tegtmeier, M., Dorn, K., Khadrawe, T.E., and Scholz, H. (2023). Octopamine integrates the status of internal energy supply into the formation of food-related memories. 10.7554/elife.88247.1.

72. Claßen, G., and Scholz, H. (2018). Octopamine Shifts the Behavioral Response From Indecision to Approach or Aversion in Drosophila melanogaster. Front Behav Neurosci 12, 131. 10.3389/fnbeh.2018.00131.

73. Scheiner, R., Steinbach, A., Claßen, G., Strudthoff, N., and Scholz, H. (2014). Octopamine indirectly affects proboscis extension response habituation in Drosophila melanogaster by controlling sucrose responsiveness. J Insect Physiol 69, 107–117. 10.1016/j.jinsphys.2014.03.011.

74. Schneider, A., Ruppert, M., Hendrich, O., Giang, T., Ogueta, M., Hampel, S., Vollbach, M., Büschges, A., and Scholz, H. (2012). Neuronal Basis of Innate Olfactory Attraction to Ethanol in Drosophila. Plos One 7, e52007. 10.1371/journal.pone.0052007.

75. Yang, Z., Yu, Y., Zhang, V., Tian, Y., Qi, W., and Wang, L. (2015). Octopamine mediates starvation-induced hyperactivity in adult Drosophila. Proc National Acad Sci 112, 5219–5224. 10.1073/pnas.1417838112.

76. Youn, H., Kirkhart, C., Chia, J., and Scott, K. (2018). A subset of octopaminergic neurons that promotes feeding initiation in Drosophila melanogaster. Plos One 13, e0198362. 10.1371/journal.pone.0198362.

77. Sayin, S., Backer, J.-F.D., Siju, K.P., Wosniack, M.E., Lewis, L.P., Frisch, L.-M., Gansen, B., Schlegel, P., Edmondson-Stait, A., Sharifi, N., et al. (2019). A Neural Circuit Arbitrates between Persistence and Withdrawal in Hungry Drosophila. Neuron 104, 544-558.e6. 10.1016/j.neuron.2019.07.028.

78. Turrel, O., Ramesh, N., Escher, M.J.F., Pooryasin, A., and Sigrist, S.J. (2022). Transient active zone remodeling in the Drosophila mushroom body supports memory. Curr Biol. 10.1016/j.cub.2022.10.017.

79. Zhang, X., Li, Q., Wang, L., Liu, Z.-J., and Zhong, Y. (2018). Active Protection: Learning-Activated Raf/MAPK Activity Protects Labile Memory from Rac1-Independent Forgetting. Neuron 98, 142-155.e4. 10.1016/j.neuron.2018.02.025.

80. Tirian, L., and Dickson, B.J. (2017). The VT GAL4, LexA, and split-GAL4 driver line collections for targeted expression in the Drosophila nervous system. bioRxiv, 198648. 10.1101/198648.

81. Stocker, R.F., Heimbeck, G., Gendre, N., and Belle J.S. de (1997). Neuroblast ablation in Drosophila P[GAL4] lines reveals origins of olfactory interneurons. J. Neurobiol. 32, 443–456. 10.1002/(sici)1097-4695(199705)32:5<443::aid-neu1>3.0.co;2-5.

82. FriggiLJGrelin, F., Coulom, H., Meller, M., Gomez, D., Hirsh, J., and Birman, S. (2003). Targeted gene expression in Drosophila dopaminergic cells using regulatory sequences from tyrosine hydroxylase. J. Neurobiol. 54, 618–627. 10.1002/neu.10185.

83. Pfeiffer, B.D., Jenett, A., Hammonds, A.S., Ngo, T.-T.B., Misra, S., Murphy, C., Scully, A., Carlson, J.W., Wan, K.H., Laverty, T.R., et al. (2008). Tools for neuroanatomy and neurogenetics in Drosophila. Proc. Natl. Acad. Sci. 105, 9715–9720. 10.1073/pnas.0803697105.

84. Mahr, A., and Aberle, H. (2006). The expression pattern of the Drosophila vesicular glutamate transporter: A marker protein for motoneurons and glutamatergic centers in the brain. Gene Expr. Patterns 6, 299–309. 10.1016/j.modgep.2005.07.006.

85. Cole, S.H., Carney, G.E., McClung, C.A., Willard, S.S., Taylor, B.J., and Hirsh, J. (2005). Two Functional but Noncomplementing Drosophila Tyrosine Decarboxylase Genes DISTINCT ROLES FOR NEURAL TYRAMINE AND OCTOPAMINE IN FEMALE FERTILITY*. J. Biol. Chem. 280, 14948– 14955. 10.1074/jbc.m414197200.

86. Sage, D., Donati, L., Soulez, F., Fortun, D., Schmit, G., Seitz, A., Guiet, R., Vonesch, C., and Unser, M. (2017). DeconvolutionLab2: An open-source software for deconvolution microscopy. Methods 115, 28–41. 10.1016/j.ymeth.2016.12.015.

87. Haase, R., Royer, L.A., Steinbach, P., Schmidt, D., Dibrov, A., Schmidt, U., Weigert, M., Maghelli, N., Tomancak, P., Jug, F., et al. (2020). CLIJ: GPU-accelerated image processing for everyone. Nat. Methods 17, 5–6. 10.1038/s41592-019-0650-1.

88. Ollion, J., Cochennec, J., Loll, F., Escudé, C., and Boudier, T. (2013). TANGO: a generic tool for high-throughput 3D image analysis for studying nuclear organization. Bioinformatics 29, 1840–1841. 10.1093/bioinformatics/btt276.

89. Kondo, S., and Ueda, R. (2013). Highly Improved Gene Targeting by Germline-Specific Cas9 Expression in Drosophila. Genetics 195, 715–721. 10.1534/genetics.113.156737.

90. Ke, M.-T., Nakai, Y., Fujimoto, S., Takayama, R., Yoshida, S., Kitajima, T.S., Sato, M., and Imai, T. (2016). Super-Resolution Mapping of Neuronal Circuitry With an Index-Optimized Clearing Agent. Cell Reports 14, 2718–2732. 10.1016/j.celrep.2016.02.057.

91. Yamagata, N., Ezaki, T., Takahashi, T., Wu, H., and Tanimoto, H. (2021). Presynaptic inhibition of dopamine neurons controls optimistic bias. Elife 10, e64907. 10.7554/elife.64907.

